# Acetylation facilitates NPM1 anchoring in the nucleolus by boosting liquid–liquid phase separation

**DOI:** 10.64898/2026.10.27.758729

**Authors:** Rafael L. Giner-Arroyo, Pablo Rivero-García, Miguel A. De la Rosa, Irene Díaz-Moreno

## Abstract

Nucleophosmin (NPM1) is a multifunctional nucleolar protein whose integration into the nucleolus relies on multivalent interactions that drive liquid–liquid phase separation (LLPS). Acetylation of Lys27 and Lys54 within the N-terminal oligomerization domain of NPM1 has been implicated in its nucleolar redistribution during the DNA damage response, but the molecular mechanisms underlying this redistribution remain unclear. Here, we show that Lys27/Lys54 acetylation facilitates NPM1 anchoring in the nucleolus under basal conditions by promoting LLPS. Despite preserving NPM1 structural integrity and oligomerization equilibrium, lysine acetylation reshapes the electrostatic surface potential of the N-terminal domain, neutralizing a predominantly basic region and exposing a previously cryptic acidic surface. This increased charge blockiness enhances the ability of NPM1 to undergo both homotypic and heterotypic LLPS. Thus, lysine acetylation acts as a modulator of NPM1 interaction landscape, promoting its integration into nucleolar condensates despite not constituting the primary determinant of its localization.

## Introduction

NPM1 is a predominantly nucleolar phosphoprotein that participates in numerous cellular processes, namely ribosome biogenesis (Ugolini *et al*., 2022; Maggi *et al*., 2008), centrosome duplication (Okuda, 2002; Wang *et al*., 2005), histone chaperoning (Okuwaki *et al*., 2001; Buzón *et al*., 2023) and DNA damage response (DDR), among others (Boulon *et al*., 2010; Lindström *et al*., 2018). In the nucleolus, NPM1 stabilizes the alternate reading frame (ARF) tumor suppressor protein and, upon DNA damage, it dissociates from ARF, which shifts out of the nucleolus and disrupts the human double minute 2 (HDM2)-mediated p53 degradation (Brady *et al*., 2004; Korgaonkar *et al*., 2005; Kurki *et al*., 2004), therefore regulating cell cycle progression. Moreover, NPM1 can also directly interact with p53, promoting its stability and transcriptional activation (Colombo *et al*., 2002; Yang *et al*., 2016). In this context, nucleus-translocated cytochrome *c* (C*c*) also interacts with NPM1 in the nucleolus to promote the release of ARF (González-Arzola *et al*., 2022; Buzón *et al*., 2023; Casado-Combreras *et al*., 2024).

NPM1 contains two structured domains: an N-terminal oligomerization domain and a C-terminal DNA binding domain, which are both linked by a central intrinsically disordered region (IDR) that alternates stretches of acidic and basic residues (Hingorani *et al*., 2000). The N-terminal domain adopts a pentameric conformation formed by five β-sheet jelly rolls (Figure 1A) (Lee *et al*., 2007; Mitrea *et al*., 2014). The multivalence provided by oligomerization, together with the charge block distribution (Yamazaki *et al*., 2022), allows NPM1 to undergo liquid-liquid phase separation (LLPS), integrating in the nucleolus by means of its self-association (Mitrea *et al*., 2018) and interaction with other nucleolar components (i.e. RNA, R-motifs) (Mitrea *et al*., 2016; Riback *et al*., 2020). Nucleolar integration through these mechanisms is highly dynamic and tightly regulated in response to cellular stress.

**Figure 1.**
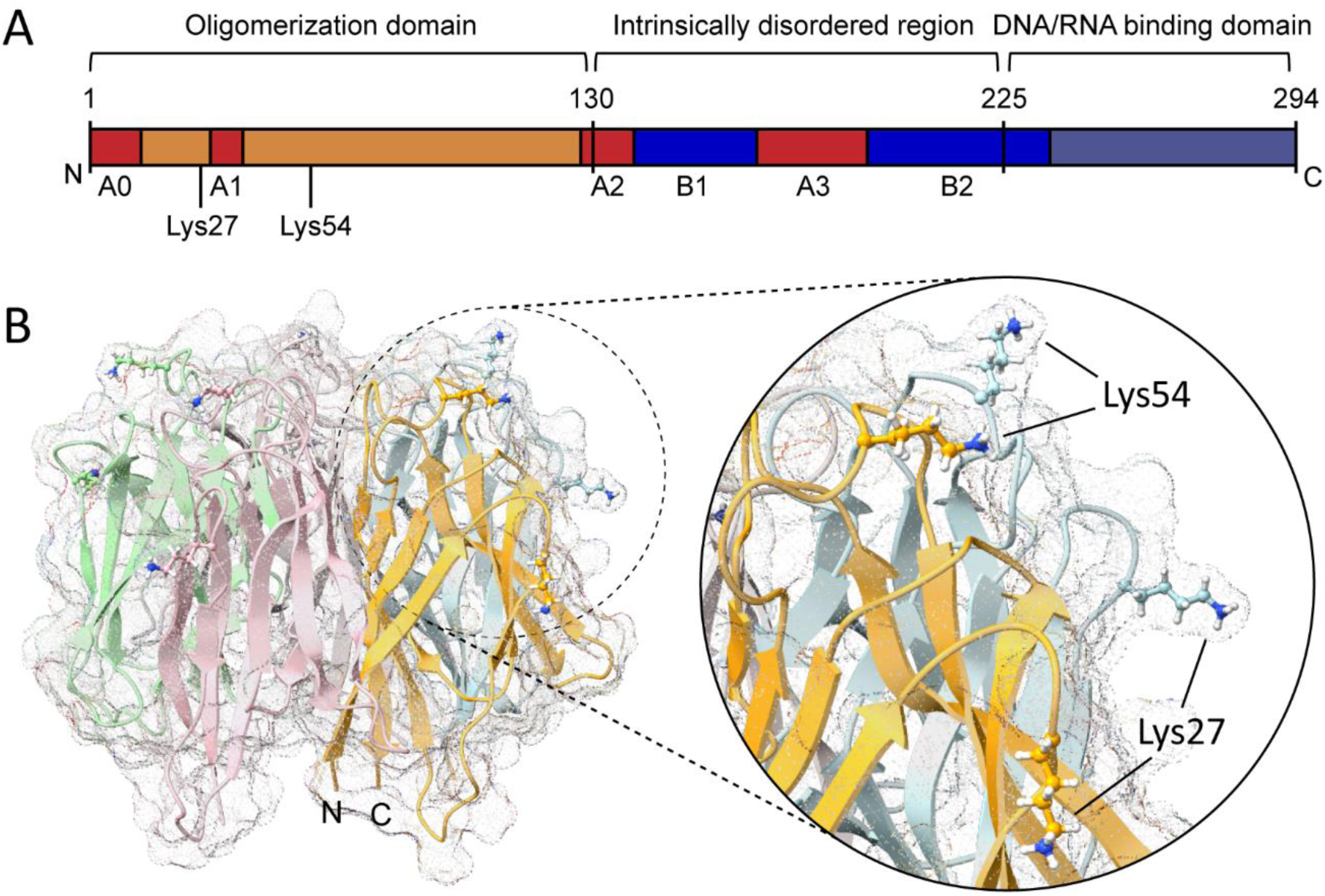
Structural organization of NPM1 and position of lysine acetylation sites. **A.** Domain organization of NPM1 comprising an N-terminal oligomerization domain, a central intrinsically disordered region and a C-terminal DNA/RNA binding domain. Acidic (A0, A1, A2 and A3) and basic (B1, B2) tracts are shown in red and blue, respectively. The position of the acetylatable residues Lys27 and Lys54 is indicated. **B.** Crystallographic structure of the N-terminal oligomerization domain (PDB: 4N8M (Mitrea *et al*., 2014)). The ribbon representation of the homopentamer is shown, with each protomer displayed in a different color. A close-up view highlights the position of the acetylatable lysines 27 and 54.

The nucleolar anchorage of NPM1 is tightly regulated by post-translational modifications (PTMs). Phosphorylation sites spanning both the N-terminal (Russo *et al*., 2021; Giner-Arroyo *et al*., 2026) and C-terminal (Sridharan *et al*., 2022; Rivero-García *et al*., 2026) domains have been shown to impair nucleolar localization. Acetylation sites Lys27 and Lys54 at the N-terminal domain of NPM1 (Figure 1B) have also been identified to be involved in the DNA damage response shuttling of the protein. Upon UV irradiation, ataxia telangiectasia mutated and Rad3-related (ATR) phosphorylates and activates the nucleolar sirtuin SIRT7, which in turn deacetylates Lys27 and Lys54 of NPM1. Subsequently, NPM1 exits the nucleolus and prevents p53 ubiquitination and degradation (Ianni *et al*., 2021). However, SIRT7 has also been proposed to promote ARF degradation by competitively disrupting the NPM1-ARF interaction, a process independent of its deacetylase activity (Kumari *et al*., 2024). Consequently, despite the identification of these acetylation sites, how acetylation at Lys27 and Lys54 influences the structural, dynamic, and functional properties of NPM1 remains largely unknown. In particular, it is unclear whether these modifications regulate NPM1 subcellular localization or, alternatively, modulate its condensation and associated functions.

In this work, we address this question by site-specifically introducing genetically encoded acetyl-lysine at positions 27 and 54 of NPM1 in mammalian cells using an efficient non-canonical amino acid incorporation system (Cohen and Arbely, 2016). By combining this cellular approach with *in silico* analysis and *in vitro* characterization of the structural dynamics, oligomerization state, and interaction properties of acetylated NPM1, we demonstrate that acetylation of Lys27 and Lys54 facilitates the anchoring of NPM1 within the nucleolus by enhancing its homotypic and heterotypic liquid–liquid phase separation. Rather than directing nucleolar localization, this PTM strengthens the intermolecular interactions that stabilize NPM1 within the biomolecular condensate, uncovering a previously unrecognized role for lysine acetylation in regulating its biomolecular condensation properties.

## Results

### Deacetylation alone does not drive UV-induced NPM1 nucleolus-to-nucleoplasm shuttling

Since UV-induced deacetylation of NPM1 has been proposed to trigger its delocalization from the nucleolus to the nucleoplasm, we assessed the impact of acetylation mimicry (Lys-to-Gln) and acetylation impairment (Lys-to-Arg) at Lys27 and Lys54 upon UV irradiation. The double lysine mutants were generated within an NPM1 fusion protein construct carrying a C-terminal *c*-myc tag (NPM1-*c*-myc) and used to transfect U2OS cells. Immunofluorescence assays revealed no differences in nucleolar localization of K27Q-K54Q and K27R-K54R NPM1-*c*-myc mutants compared to WT NPM1-*c*-myc in non-stressed cells and, regardless of lysine mutations, all variants similarly displayed a more nucleoplasmic distribution in UV treated cells. Simultaneous immunostaining of NPM1 and *c*-myc tag allowed to differentiate between total and transfected NPM1 (Figure 2).

**Figure 2.**
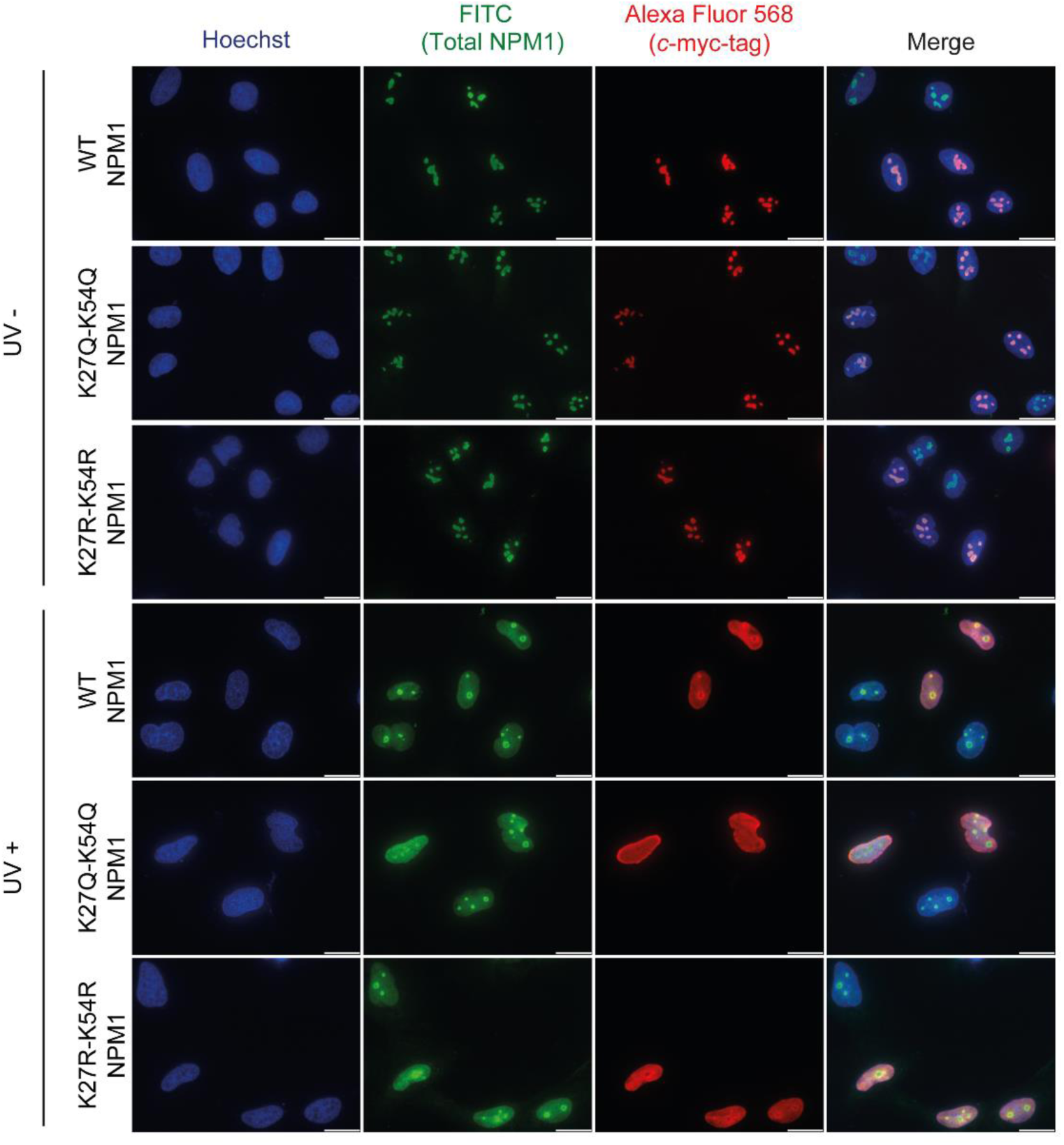
Subcellular localization of NPM1 acetyl-mimetic and acetylation-impaired mutants in the presence and absence of UV stress. Immunofluorescence analysis of U2OS cells transfected with wild type (WT), double acetyl-mimetic (K27Q-K54Q) or acetylation-impaired (K27R-K54R) NPM1 variants upon UV genotoxic stress using anti-NPM1 and anti-*c*-myc tag antibodies. When indicated, cells were treated with 80 J m^-2^ UV light. Scale bars, 20 μm.

To unequivocally address the impact of acetylation in NPM1 cellular localization, we resorted to the incorporation of genetically-encoded acetyl-lysine (AcK) in positions 27 and 54 of NPM1 using an Amber suppression one-plasmid system, designed to optimize protein expression in cultured mammalian cells (Cohen and Arbely, 2016). The co-expression in U2OS cells of NPM1-*c*-myc K27Amber, K54Amber or K27Amber-K54Amber and the orthogonal aminoacyl tRNA synthetase/tRNA pair allowed for the expression of acetylated variants without the need of co-transfection. Incorporation of AcK was confirmed through immunostaining of the C-terminal *c-*myc-tag, which was only detected when AcK non-canonical amino acid was present in the culture medium (Figures 3 and S1). Cells were treated with 1 mM nicotinamide to inhibit sirtuin-mediated deacetylation and thereby maintaining NPM1 in an acetylated state. Similar to WT NPM1-*c*-myc, single and double acetylated NPM1-*c*-myc variants displayed nucleolar localization in untreated cells and redistributed to the nucleoplasm upon UV irradiation (Figure 3). Acetylation levels remained detectable 24 h after transfection in untreated cells, as confirmed by anti-*c*-myc-tag immunoprecipitation of the double acetylated *c*-myc-tagged protein followed by anti-acetyl-lysine Western Blot (Figure S2). Altogether, these results show that doubly acetylated NPM1 predominantly localizes to the nucleolus in the absence of genotoxic stress.

**Figure 3.**
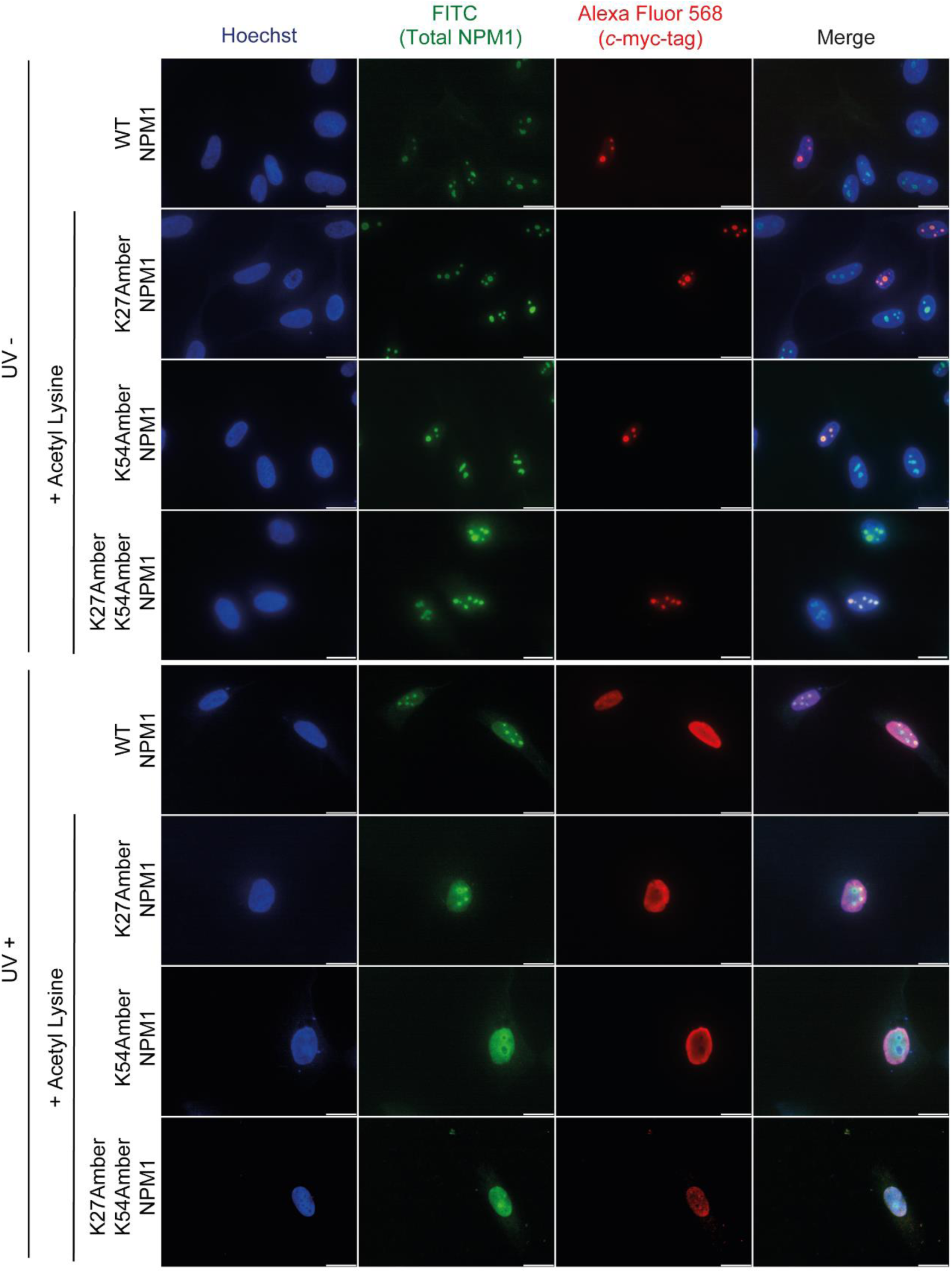
Subcellular localization of NPM1 acetylated variants in the presence and absence of UV stress. Immunofluorescence staining of U2OS cells transfected with NPM1 Amber variants for the incorporation of genetically encoded acetyl-lysine in positions 27, 54 or both. When indicated, cells were irradiated with 80 J m^-2^ UV light. Total and transfected NPM1 were stained using anti-NPM1 and anti-myc tag antibodies, respectively. Scale bars, 20 μm.

### Acetylation enhances the acidic nature of NPM1 oligomerization domain while preserving its structural integrity, Lys27/Lys54 contact network, and oligomeric equilibrium

The integration of NPM1 into the nucleolus relies on its capacity of self-association (Mitrea *et al*., 2018) and interaction with other nucleolar components (Mitrea *et al*., 2016; Riback *et al*., 2020). These are highly dependent on its oligomeric structural architecture and charged tracts distribution (Figure 1A), which enables the establishment of multivalent interactions (Mitrea *et al*., 2018; Giner-Arroyo *et al*., 2026). Since both Lys27 and Lys54 are located in the oligomerization domain of NPM1 (Figure 1B), we examined the electrostatic surface potential (ESP) of NPM1 oligomerization domain in WT and doubly acetylated (K27AcK-K54AcK) variants. Notably, acetylation of Lys27 and Lys54 not only neutralized their positive charges, but also substantially remodeled the local electrostatic potential, shifting the surrounding surface from predominantly basic to predominantly acidic (Figure 4).

**Figure 4.**
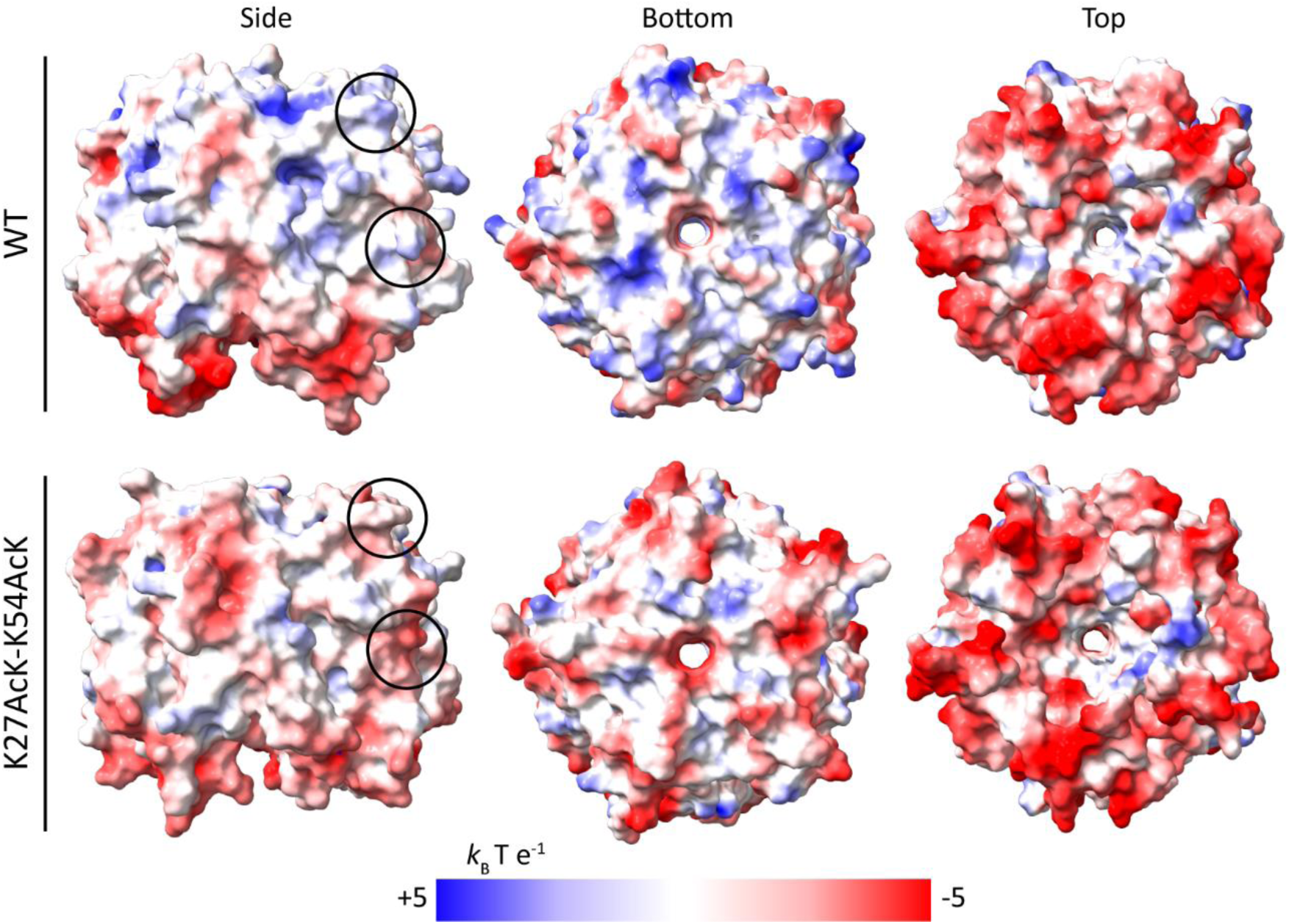
**Electrostatic surface potential (ESP) of NPM1 oligomerization domain in WT and doubly-acetylated (K27AcK-K54AcK) variants**. ESPs were calculated at 100 mM ionic strength using APBS (Jurrus *et al*., 2018) on representative structures obtained on the molecular dynamics simulations. The position of the two lysines in one of the protomers is indicated by circles.

We also studied the oligomerization equilibrium of acetyl-mimetic variants. The pentameric structure of NPM1 has been shown to persist through SDS-PAGE, even in the presence of the 1% SDS of the sample buffer, provided that samples are not boiled (Okuwaki *et al*., 2012; Okuwaki *et al*., 2023). Nonetheless, this oligomeric equilibrium has previously been reported to be affected by several PTMs including serine/threonine (Mitrea *et al*., 2014) and tyrosine phosphorylation (Giner-Arroyo *et al*., 2026). To evaluate the impact of acetylation on NPM1 oligomer stability, acetyl-mimetic variants were incubated at increasing temperatures prior to SDS–PAGE analysis. All variants exhibited comparable monomer–oligomer distributions across the temperature range tested, remaining predominantly pentameric in the absence of heating and almost exclusively monomeric after boiling at 95 °C. The multi-banded oligomeric pattern likely reflects the presence of higher-order assemblies, including decamers, as previously reported (Lee *et al*., 2007; Sakashita, 2018) (Figure 5). Collectively, these results indicate that acetylation of Lys27 and Lys54 does not significantly affect the stability of NPM1 oligomers.

**Figure 5.**
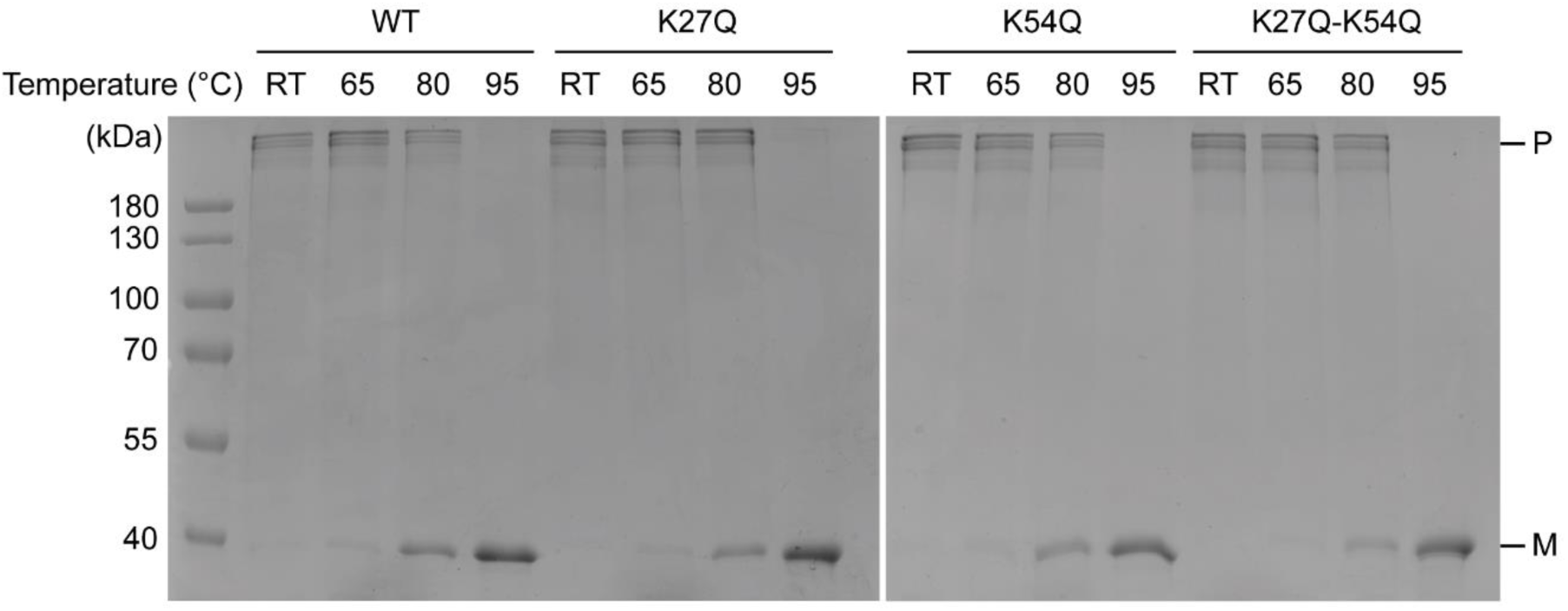
Temperature-dependent stability of NPM1 oligomers in Lys27 and Lys54 acetyl-mimetic mutants. SDS-PAGE analysis of the oligomeric distribution of WT and acetyl-mimetic NPM1 single (K27Q, K54Q) and double (K27Q-K54Q) mutant variants after 5 min heating at RT, 65, 80 or 95 °C. P and M indicate bands corresponding to pentamers and monomers, respectively.

To investigate whether acetylation (AcK), acetyl-mimetic (Gln), or acetylation-impaired (Arg) substitutions affect the stability of the NPM1 oligomerization domain and the contact network surrounding Lys27 and Lys54, we performed molecular dynamics (MD) simulations. Three independent 1-µs simulations were carried out for each variant using the NPM1 oligomerization domain (PDB: 4N8M) as the starting structure. Structural stability was assessed by analyzing the root mean square deviation (RMSD), radius of gyration (RoG), and residue contact frequencies.

The RMSD and RoG converged to similar values for all variants, suggesting that the overall structural integrity of NPM1 is preserved across different substitutions (Figures 6A, 6B, 6C). Moreover, the identity of the main residues participating in contacts within the pentamer with Lys27 (Glu22, Leu23, Lys24) and Lys54 (His58, Ser106, Gly107) remained constant among all variants. However, Lys27 acetylation showed an increase on the contact frequency with its partner residues, which was also observed with the Gln substitution in both single and double mutant, validating its acetyl-mimetic behavior (Figures 6D, 6E). Nonetheless, acetylation could also influence the contacts between the oligomerization domain and the intrinsically disordered acidic central region of NPM1. In this context, the shift in the ESP of the oligomerization domain upon lysine acetylation, resulting in a predominantly acidic surface, is expected to modulate such contacts and, eventually, the propensity of NPM1 to undergo LLPS by reshaping the protein’s charge blockiness (Yamazaki *et al*., 2022).

**Figure 6.**
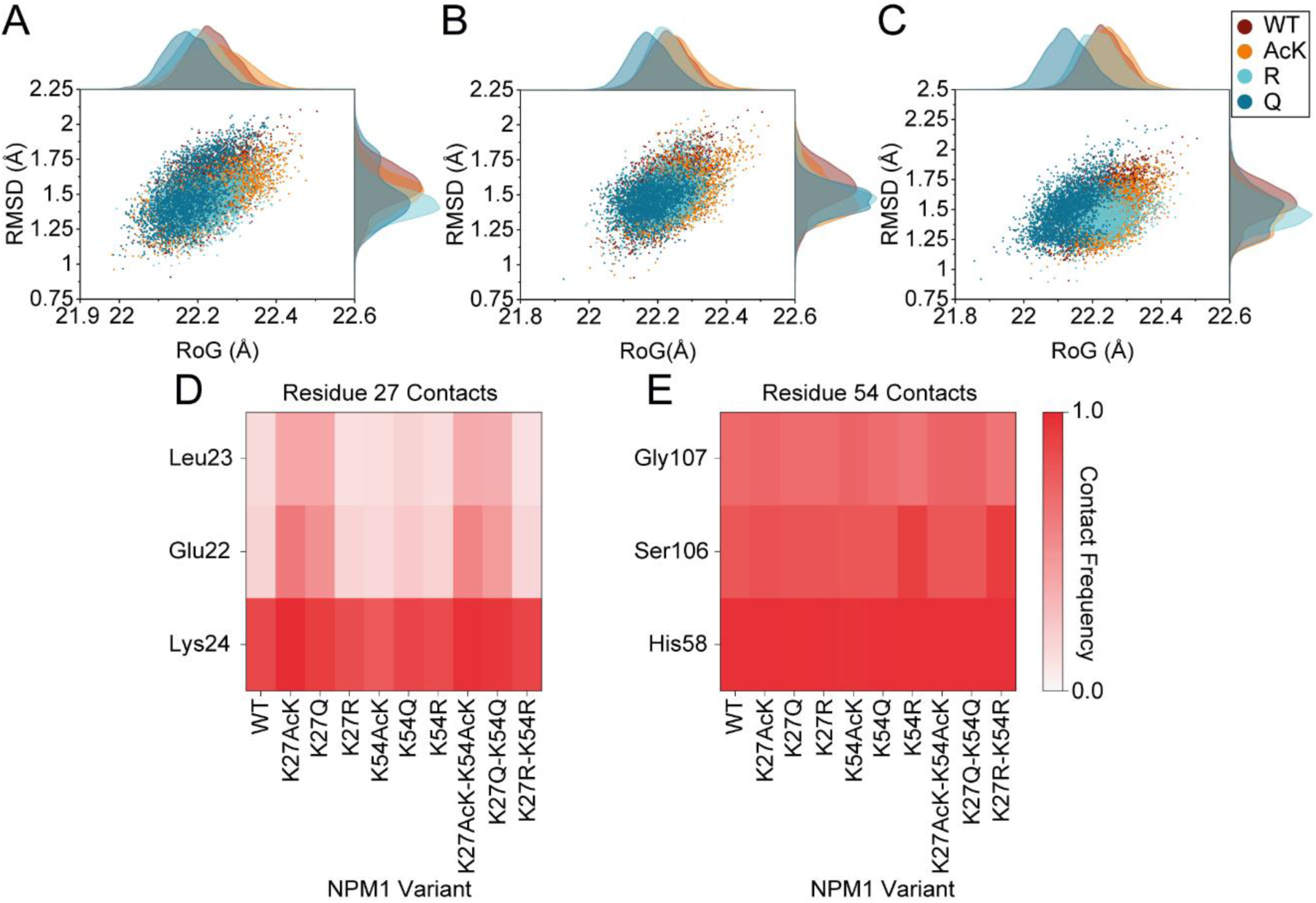
Molecular Dynamics analysis of the N-terminal oligomerization domain in NPM1 acetylation variants. A, B and. **C**. RMSD and RoG correlation of NPM1 variants mutated in positions 27 (**A**), 54 (**B**) or both (**C**) during 1-μs MD simulations. **D and E.** Frequency heatmaps of the most frequent contacts of residues in position 27 (**D**) and 54 (**E**) in the simulations of the NPM1 variants. The fraction of the 1-µs simulation during which the residue pairs remain in contact is represented by a color scale from 0 to 1.

### Acetylation enhances NPM1 liquid-liquid phase separation

NPM1 incorporation into the nucleolus is closely linked to its ability to undergo LLPS by both self-association and interaction with other nucleolar components. Homotypic LLPS relies on the electrostatic complementarity of basic and acidic tracts in NPM1 IDR and oligomerization domain (Mitrea *et al*., 2018). Heterotypic NPM1 LLPS—particularly with its partners p19ARF and C*c*, both of which participate in the NPM1-dependent regulation of p53 (Brady *et al*., 2004; González-Arzola *et al*., 2022)—is also driven by electrostatic interactions between the multivalent basic motifs and acidic tracts of NPM1 (Mitrea *et al*., 2016). In both cases, multivalency arising from NPM1 pentamerization is essential for LLPS.

While NPM1 oligomerization is not significantly impacted by acetylation, as inferred from SDS-PAGE and molecular dynamics simulations, we suggest that the neutralization of lysine positive charges upon acetylation affects electrostatic contacts. Thus, to gain more insights into the contribution of NPM1 acetylation to its nucleolar integration, we performed *in vitro* phase separation assays on NPM1 acetyl-mimetic mutants. Homotypic LLPS tendency of the NPM1 variants was addressed by the addition of the crowding agent Ficoll across several protein concentrations (Figure 7A). All variants similarly formed liquid droplets with comparable morphologies, but acetyl-mimetic variants displayed a modest, yet generalized increase as revealed by the quantification (Figure 7B). This increase may result from the acetylation-induced reshaping of the ESP, whereby acidification of the electrostatic surface surrounding Lys27 and Lys54 (Figure 3) expands the contribution of the N-terminal oligomerization domain to homotypic LLPS beyond its acidic A0 and A1, increasing its electrostatic complementarity with the basic tracts that mediate homotypic interactions.

**Figure 7.**
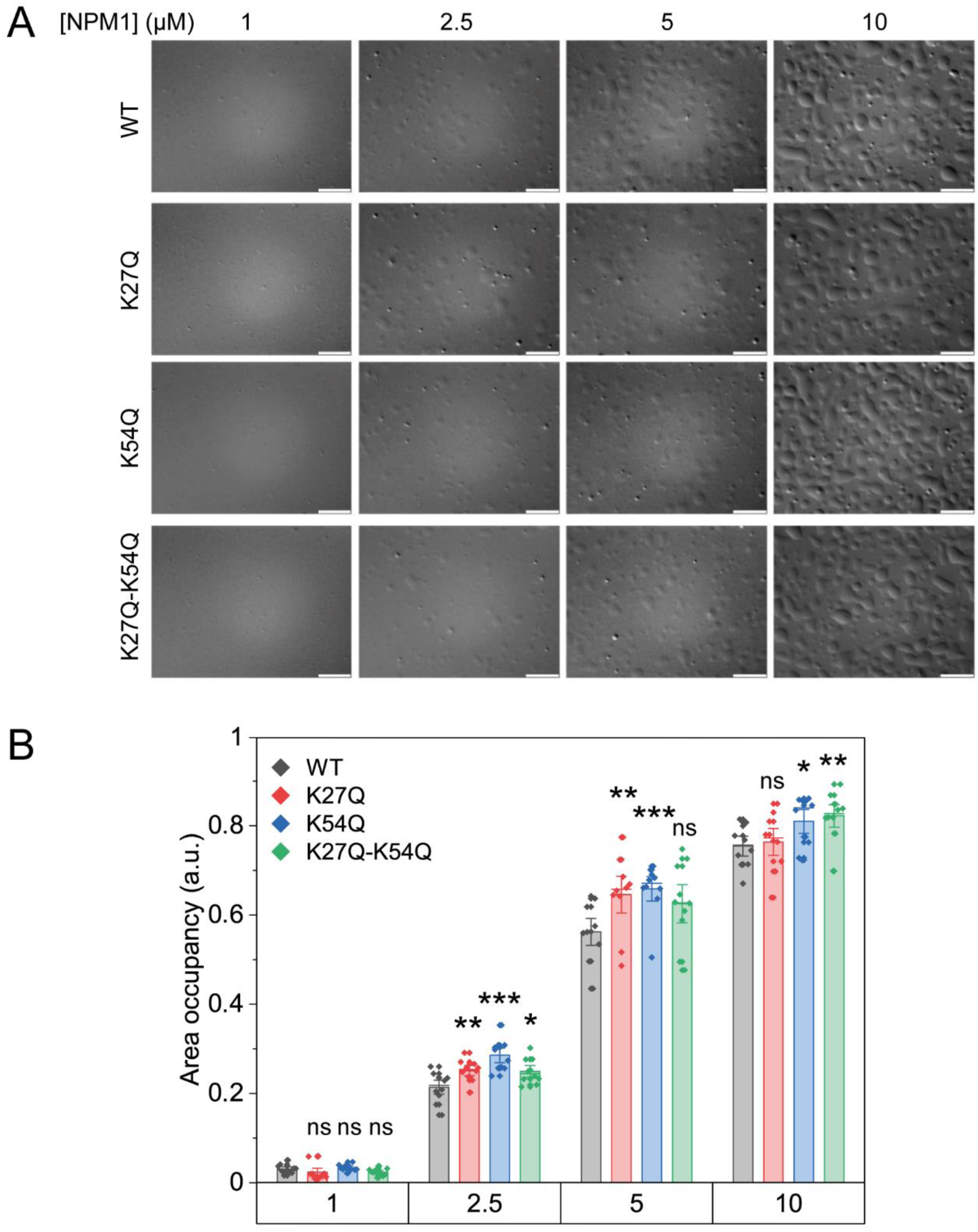
Effect of acetyl-mimetic substitutions on NPM1 homotypic phase separation. **A.** DIC microscopy images of homotypic protein condensates formed at different concentrations of WT and acetyl-mimetic NPM1 mutants in the presence of 15% Ficoll PM70. Scale bars, 10 µm. **B.** Quantification of the area occupancy of condensates in the microscopy fields. Bars represent the mean and error bars indicate ± one standard deviation. Statistically significant differences relative to WT are indicated: *P* < 0.05 (*), *P* < 0.01 (**), *P* < 0.005 (***), not significant (ns).

Furthermore, heterotypic droplets formed between NPM1 and p19ARF were more abundant with acetyl-mimetic single mutants at some of the concentration ratios tested and showed a consistent increase across all ratios in the double mutant (Figures 8A, 8B). By contrast, NPM1–C*c* readily formed abundant heterotypic droplets across all variants, with only minor differences in droplet abundance between them (Figures 9A, 9B). Interestingly, neither ARF nor C*c* appears to make direct contacts with the acetylated lysines or with the residues surrounding these sites in the available NPM1–partner structures (PDB ID: 5EHD; González-Arzola *et al*., 2022; Gibbs *et al*., 2024). Thus, the differential response of ARF and C*c* to NPM1 acetylation is unlikely to result from direct interactions with the modified residues themselves, but may rather reflect differences in the physicochemical properties of the basic residues that mediate their interactions with NPM1. Whereas ARF is predominantly arginine-rich (R-rich), C*c* is primarily lysine-rich (K-rich). As discussed above, acetylation reshapes the electrostatic surface of the NPM1 pentamer by generating additional acidic character around Lys27 and Lys54. The R-rich regions of ARF may be particularly sensitive to this redistribution of charge, owing to the ability of arginine to establish multiple and more diverse interactions with acidic surfaces, whereas the K-rich sequences of C*c* may respond less strongly to these changes. In particular, the positive charge is largely localized on the terminal amino group in Lys, whereas it is delocalized over the planar guanidinium group in Arg (Gobbi and Frenking, 1993). This delocalization endows Arg with distinctive interaction properties, including a propensity to engage in cation–π interactions with aromatic residues such as Tyr and Phe, which is not observed to the same extent for Lys. In addition, the guanidinium group provides multiple hydrogen-bond donors and can establish highly directional, bidentate interactions with acidic residues (Wang *et al*., 2018; Ukmar-Godec *et al*., 2019; Casado-Combreras *et al*., 2025). By contrast, although the primary amine of Lys is less sterically constrained, it is generally limited to a single optimally oriented hydrogen bond, with additional interactions requiring deviations from ideal geometry. This difference could therefore contribute to the enhanced heterotypic LLPS observed between acetylated NPM1 and ARF, while having a more limited effect on NPM1–C*c* phase separation.

**Figure 8.**
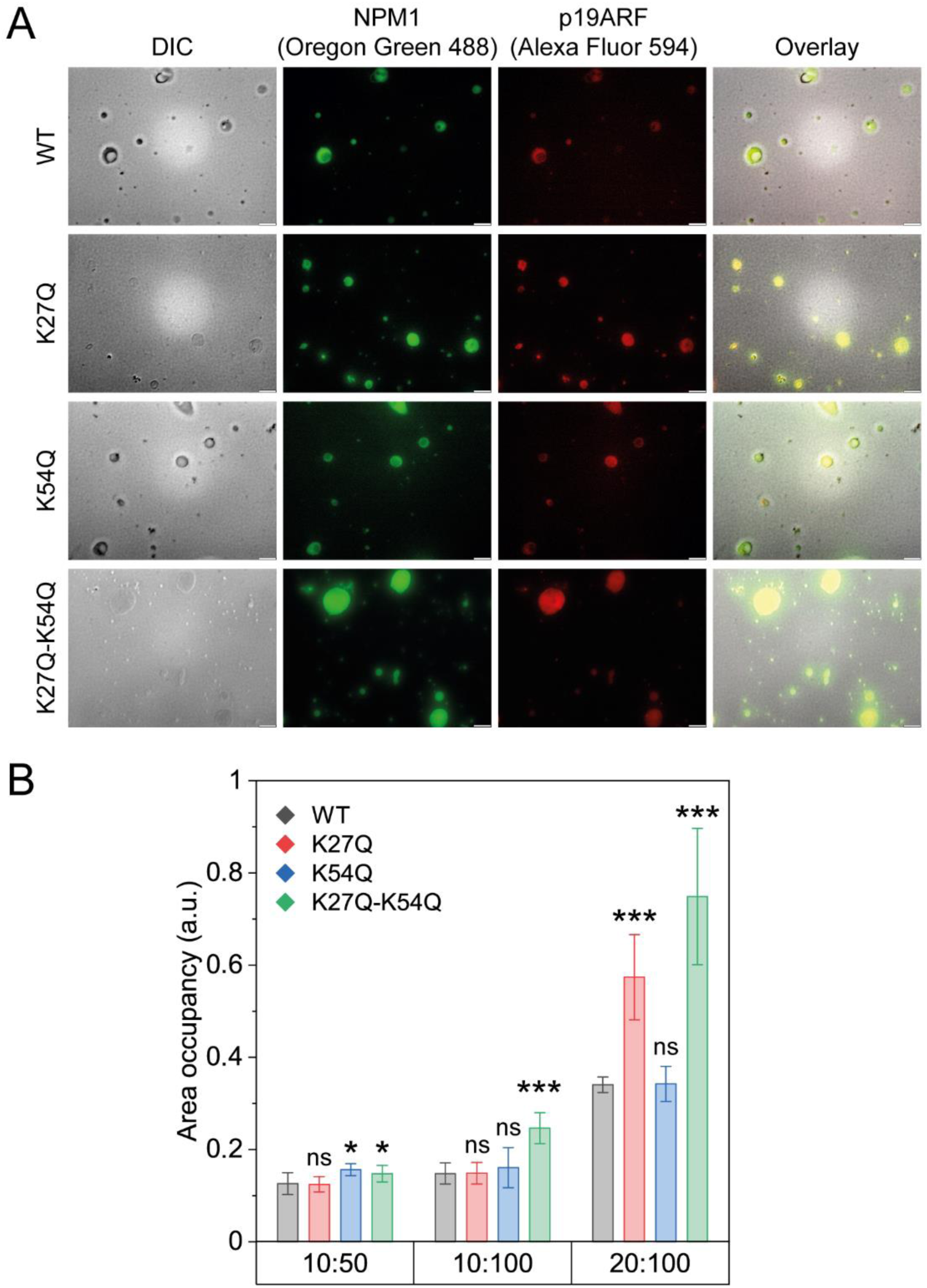
Effect of acetyl-mimetic substitutions on NPM1 heterotypic phase separation with ARF. **A.** WT and acetyl-mimetic NPM1 mutants were incubated with synthetic p19ARF_(1-37)_ peptide at different concentration ratios. Representative microscopy images of condensates formed at an NPM1:p19ARF_(1-37)_ ratio of 20:100 μM are shown by DIC, Oregon Green 488 fluorescence from labelled NPM1 and Alexa Fluor 594 from labelled p19ARF_(1-37)_. Scale bars, 10 µm. **B.** Quantification of the area occupancy of condensates in the microscopy fields. Bars represent the mean and error bars indicate ± one standard deviation. Statistically significant differences relative to WT are indicated: *P* < 0.05 (*), *P* < 0.01 (**), *P* < 0.005 (***), not significant (ns).

**Figure 9.**
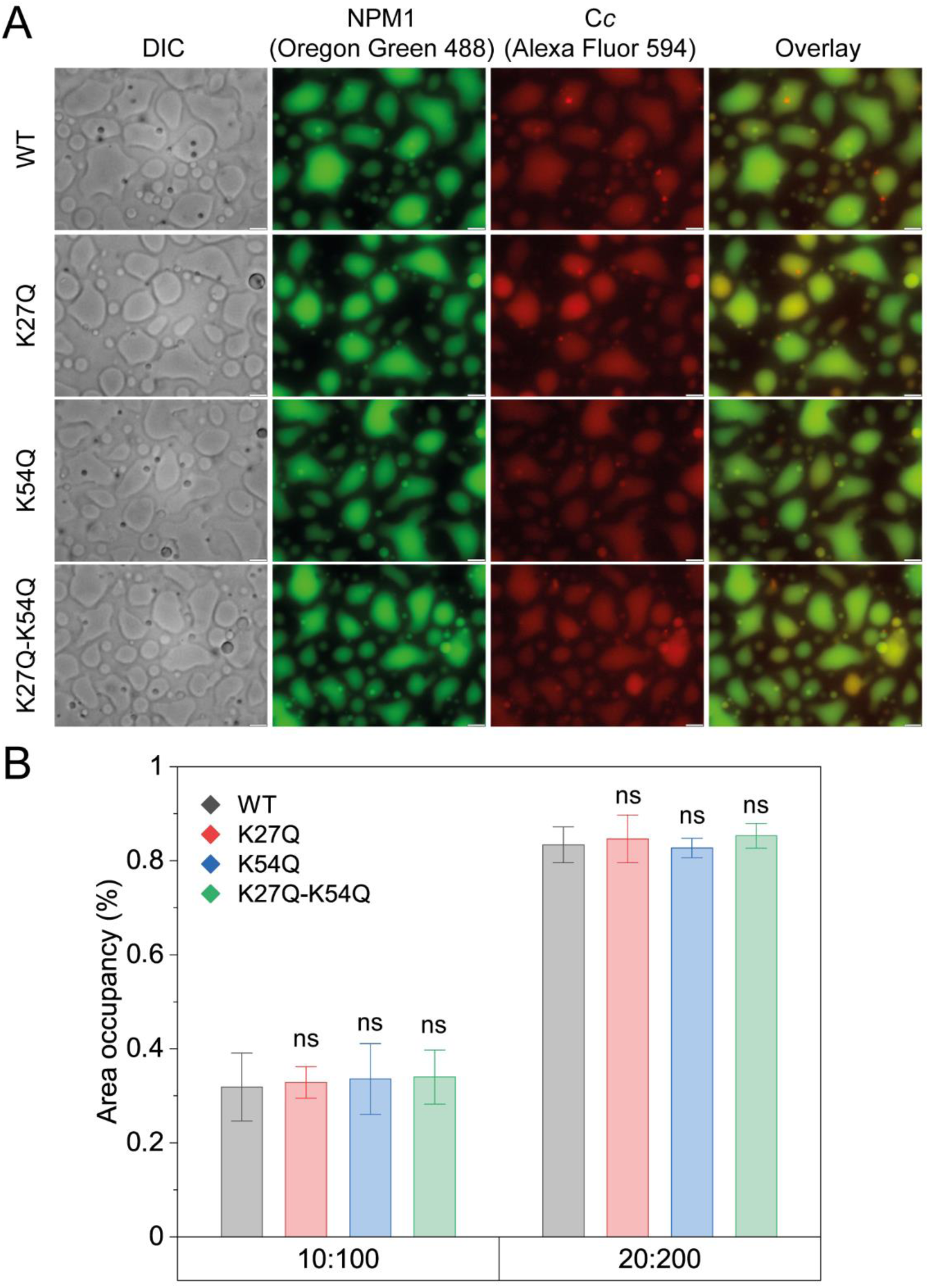
Effect of acetyl-mimetic substitutions on NPM1 heterotypic phase separation with C*c*. **A.** WT and acetyl-mimetic NPM1 mutants were incubated with C*c* at different concentration ratios. Representative microscopy images of condensates formed at an NPM1:C*c* ratio of 20:200 μM are shown by DIC, Oregon Green 488 fluorescence from labelled NPM1 and Alexa Fluor 594 from labelled C*c*. Scale bars, 10 µm. **B.** Quantification of the area occupancy of condensates in the microscopy fields. Bars represent the mean and error bars indicate ± one standard deviation. Statistically significant differences relative to WT are indicated: *P* < 0.05 (*), *P* < 0.01 (**), *P* < 0.005 (***), not significant (ns).

This difference in the interaction potential of R-and K-rich sequences may influence not only the extent of heterotypic LLPS but also the material properties of the resulting condensates. Notably, the condensates formed with ARF displayed limited coalescence and reduced wetting compared with those formed with C*c*, consistent with a denser intermolecular interaction network mediated by the multivalent R-rich motifs, as explained before (González-Arzola *et al*., 2022). On the contrary, wetting reflects the balance between cohesive forces within the condensate and adhesion to the solid surface (Wang and Shi, 2025). Thus, the weaker and less diverse interactions mediated by the K-rich C*c* may lead to reduced condensate cohesion and increased fluidity, potentially accounting for the distinct morphology and wetting behavior observed for NPM1–C*c* condensates.

## Discussion

Deacetylation of NPM1 Lys27 and Lys54 by SIRT7 has been proposed to be directly involved in the stabilization of p53 following the UV-induced DDR by promoting the redistribution of NPM1 from the nucleolus to the nucleoplasm (Ianni *et al*., 2021). However, more recent work has shown that SIRT7 enzymatic activity is not necessary for modulation of the NPM1/ARF/p53 axis and instead suggests that SIRT7 acts as a competitive inhibitor that prevents NPM1-ARF binding (Kumari *et al*., 2024). Here, we demonstrate that Lys27 and Lys54 acetylation facilitates the anchoring of NPM1 in the nucleolus by enhancing NPM1 phase separation through homo- and heterotypic contacts with physiological partners such as ARF.

Immunofluorescence analysis revealed that NPM1 remained predominantly nucleolar even when acetylation was prevented by the K27R-K54R substitutions, indicating that the deacetylation at these sites is not sufficient to trigger NPM1 release from the nucleolus. Moreover, UV irradiation induced NPM1 redistribution to the nucleoplasm regardless of the acetylation status of Lys27 and Lys54, as observed for the acetylation-impaired (K27R-K54R), acetyl-mimetic (K27Q-K54Q), and genetically acetylated (K27AcK-K54AcK) variants.

Notably, both Lys27 and Lys54 face towards the surface of the N-terminal oligomerization domain and are located considerably far from the interface between monomers (Figure 1B). Both the oligomerization state of NPM1 and the stability of the N-end domain were not significantly affected by lysine acetylation, as supported by our MD simulation and SDS-PAGE. Instead of altering NPM1 structure, acetylation primarily reshaped its electrostatic surface potential and, consequently, its interaction landscape, enhancing both homotypic interactions and heterotypic interactions with ARF.

Recent studies have shown that phosphorylation within the intrinsically disordered central region of NPM1 can modulate its charge blockiness by reducing the net charge of basic patches, thereby weakening the electrostatic interactions that drive LLPS (Yamazaki *et al*., 2022). Building on this concept of charge block-driven LLPS, we propose that lysine acetylation in the N-terminal oligomerization domain of NPM1 acts through a complementary mechanism. By neutralizing basic charges at Lys27 and Lys54, acetylation reshapes the spatial distribution of charges across the NPM1 pentamer, enhancing the acidic character of this domain and extending its contribution beyond the canonical acidic A0 and A1 tracts. Rather than creating a new interaction surface, acetylation may therefore unmask a pre-existing but cryptic electrostatic feature of the NPM1 pentamer, generating a latent phase-separation capacity that becomes more readily engaged upon acetylation. The resulting increase in charge blockiness may enhance electrostatic complementarity with basic regions of neighboring NPM1 molecules, thereby promoting homotypic LLPS, while also increasing the ability of the acetylated NPM1 surface to engage basic motifs in heterotypic partners. Thus, Lys27 and Lys54 acetylation may act as drivers of NPM1 phase separation by reorganizing charge blocks rather than simply altering the overall net charge.

The differential response of NPM1 to acetylation in its interactions with ARF and C*c* further supports this model. Whereas NPM1 acetylation enhanced LLPS with the R-rich protein ARF, no significant effect was observed with the K-rich C*c*. This difference is unlikely to result from direct recognition of the acetylated residues, as neither partner appears to engage directly with Lys27, Lys54, or their surrounding residues in the available structural models. Instead, it may arise from the distinct chemical properties of their basic residues. The guanidinium group of Arg can establish multiple, highly directional interactions with acidic surfaces, whereas the terminal amino group of Lys has more limited interaction valence. Consequently, R-rich motifs in ARF may be better positioned to exploit the additional acidic character generated on the NPM1 surface upon acetylation, whereas the K-rich motifs of C*c* may be less sensitive to this redistribution of charge.

Such differential sensitivity to NPM1 acetylation of ARF and C*c* may have functional consequences for the composition and organization of nucleolar condensates and, consequently, for the recruitment and activity of NPM1 partners involved in stress, during DDR and p53 signaling. Our results may favor the incorporation of ARF into NPM1-containing nucleolar condensates, potentially contributing to its stable nucleolar localization under homeostatic conditions. Conversely, deacetylation upon genotoxic stress could weaken these interactions and facilitate the competition of C*c* following its nuclear translocation from mitochondria in response to DNA damage (Nolin *et al*., 2016; González-Arzola *et al*., 2022). This scenario would be consistent with the mechanism we previously proposed, whereby C*c* reaches the nucleolus and competes with ARF for NPM1 binding, promoting the release of ARF to the nucleoplasm (González-Arzola *et al*. 2022).

Accordingly, lysine acetylation facilitates the anchoring of NPM1 within the nucleolus by enhancing its LLPS properties, whereas UV-induced nucleolar exclusion requires additional regulatory mechanisms beyond deacetylation. In contrast to acetylation, other PTMs have been shown to exert a much stronger impact on the phase separation behavior of NPM1. By disrupting monomer–monomer contacts and promoting unfolding of the N-terminal domain, phosphorylation at Tyr17 and Tyr67 destabilizes the pentamer, severely impairing both homotypic and heterotypic LLPS (Giner-Arroyo *et al*., 2026). These structural changes ultimately drive NPM1 translocation from the nucleolus to the nucleoplasm, leading to p53 activation and the maintenance of genomic stability (Russo *et al*., 2021). Likewise, phosphorylation of Ser254, Ser260, and Tyr271 within the C-terminal nucleic acid-binding domain also promotes NPM1 nucleoplasmic relocalization by weakening nucleic acid binding and disrupting charge block-driven homotypic LLPS (Sridharan *et al*., 2022; Rivero-García *et al*., 2026).

The concomitant occurrence of NPM1 Lys27 and Lys54 deacetylation and the phosphorylation events following UV-induced genotoxic stress raises the possibility of a synergistic mechanism. While not determinant on its own, NPM1 deacetylation may contribute to debilitate some selected interactions, which could result in NPM1 being more weakly retained in the nucleolus. This would facilitate subsequent post-translational modifications, e.g. phosphorylation, that progressively shift the equilibrium toward nucleolar exclusion, ultimately enabling the complete activation of the DDR. The high solvent accessibility of Lys27 and Lys54 (Figure S3) is compatible with their deacetylation occurring as early steps of the modification sequence.

In conclusion, our findings identify Lys27/Lys54 acetylation as a modulator of NPM1 phase separation rather than a primary determinant of its nucleolar localization during the UV-induced DNA damage response. Acetylation at these sites preserves the structural integrity of the NPM1 oligomerization domain while subtly strengthening the homotypic and heterotypic interactions that drive liquid–liquid phase separation, thereby promoting the retention of NPM1 within the nucleolus.

## Materials and Methods

### DNA constructs

pBudCE4.1-based plasmids for incorporation of acetyl-lysine through AMBER suppression technique in mammalian cells were kindly provided by Eyal and co-workers (Cohen and Arbely, 2016). DNA coding sequence for NPM1 fused with a C-terminal *c*-myc-tag were inserted under the regulation of the EF1α promoter in this system by InFusion HD EcoDry^TM^ Cloning Kit (639690, Takara) following the obtention of the insert from pcDNA3.1 NPM1-*c*-myc and the template from pBud-AcKRS1-eGFP150Amber-4×tRNA by PCR amplification. pcDNA3.1 plasmid containing *c*-myc-tagged WT and double Lys-to-Gln/Arg NPM1 mutants were used to transfect U2OS cells. Mutagenic PCR was carried out to switch Lys27 and Lys54 codons to Arg, Gln or Amber codons in the different plasmids using the primers detailed in Table S1. The plasmids used in this work are listed in Table S2.

For heterologous protein expression in *E. coli*, pET28(+) bacterial expression vector containing NPM1 sequence fused with an N-terminal His-tag was already available at our group (González-Arzola *et al*., 2022). pEVOL plasmid was also used for the incorporation of AcK through Amber suppression technique. For the recombinant expression of C*c*, a pBTR1 vector which also contained the *CYC3* gene, encoding the yeast heme lyase required for the proper folding of C*c* when expressed in bacterial cell cultures was used (Olteanu *et al*., 2003; Moreno-Beltrán *et al*., 2015).

### Cell cultures and *in-cell* acetyl-lysine incorporation assay

U20S cells (100,000 per well) were grown for 24 h on 20 mm glass coverslips on 24-well plates containing 500 μL Dulbecco’s modified Eagle’s medium (DMEM) (Sigma-Aldrich) supplemented with 10% heat-inactivated fetal bovine serum (FBS) (Sigma-Aldrich), 2 mM L-glutamine (Sigma-Aldrich), 100 U mL^-1^ streptomycin (Sigma-Aldrich) and 100 μg mL^-1^ penicillin (Sigma-Aldrich) at 37 °C in a humidified atmosphere of 5% CO_2_.

Cells were then transfected with the Lipofectamine^TM^ 3000 Transfection Reagent (Invitrogen) following the manufacturer’s instructions. For each well, 1.5 μL of lipofectamine was added to 25 μL Opti-MEM^TM^ reduced serum-free media and vortexed for 2-3 s. Separately, 1 μg of plasmid DNA and 1 μL P300 Reagent were added to 25 μL of Opti-MEM^TM^ medium and mixed. Both solutions were mixed and incubated for 15 min at room temperature and the 50 μL mix was added dropwise to the cultured cells. For acetyl-lysine incorporation, the medium was supplemented with 5 mM Nε-acetyl-L-lysine (J64139.06, Alfa Aesar™) and 1 mM nicotinamide (N0636, Sigma-Aldrich). When indicated, the medium was replaced for PBS and cells were irradiated with 80 J m^-2^ UV light 24 h after transfection to induce DNA damage, and left incubating for another 17 h.

## Immunofluorescence

The growth medium was discarded, and the cells washed with phosphate buffer saline (PBS) and fixed for 15 min at room temperature in PBS containing 4% paraformaldehyde. After washing in PBS, they were permeabilized in PBS supplemented with 0.1 % Triton X-100 for 10 min at room temperature. After another washing step, cells were treated with a blocking solution of 5% bovine serum albumin (BSA, Sigma Aldrich) in PBS for 30 min at room temperature, and then incubated with mouse anti-*c*-myc tag antibody (1:500) (05-724, Sigma Aldrich) and rabbit anti-NPM antibody (1:400) (Ab183340, Abcam) for 1 h. Coverslips were washed again with PBS and probed with secondary antibodies anti-mouse IgG-Alexa Fluor 568 (1:400) (Ab175473, Abcam) and anti-rabbit IgG–FITC (1:400) (F9887, Sigma-Aldrich) for 1 h.

Nuclei were stained incubating the cells with 200 μg mL^-1^ Hoechst dye (Sigma-Aldrich) for 10 min after probing with the secondary antibodies. Leftover dye was washed with PBS. Coverslips were then immersed in ethanol for 2 min and left to dry afterwards. They were mounted for visualization using *n*-propyl gallate and sealed with nail polish. Cell imaging was carried out in a Leica CTR 6000 microscope, using a 40x objective and an ORCA-ER camera (Hamamatsu). Differential interference contrast (DIC) and fluorescence filters FITCL5 (excitation band-pass, 480/40; emission band-pass, 527/30) and Tx2 (excitation, 560/40 nm; emission, 645/75 nm) were used to visualize FITC and Alexa Fluor 568, respectively. Images were processed and analyzed with the LAS X (Leica Microsystems) and the FIJI (ImageJ) software packages.

## Immunoprecipitation and Western Blot

U2OS cells were grown to 70% confluence in a T75 flask with 15 mL of complete DMEM (supplemented as described above). Cells were transfected with the pBudCE4.1-based plasmid codifying for K27Amber-K54Amber NPM1 using Lipofectamine^TM^ 3000 Transfection Reagent (Invitrogen). 60 μL of Lipofectamine^TM^ 3000 were added to 1000 μL of Opti-MEM^TM^ reduced serum–free medium and vortexed for 2–3 s. Separately, 40 μg of plasmid DNA and 80 μL of P3000^TM^ Reagent were added to 1000 μL of Opti-MEM^TM^ medium and mixed thoroughly. Both solutions were combined and incubated for 15 min at room temperature. The resulting 2 mL transfection mix was added dropwise to the cultured cells and the medium was supplemented with 5 mM acetyl-lysine and 5 mM nicotinamide.

After 24 h, cells were lysed in 1 mL of a buffer containing 10 mM Tris-HCl pH 7.4, 150 mM NaCl, 1 mM EDTA, 1 mM EGTA, 5 mM nicotinamide, cOmplete^TM^ EDTA-free Protease Inhibitor Cocktail (Roche), 1% Triton X-100 and 0.5% NP-40, in constant agitation for 30 min at 4 °C. Immunoprecipitation of the recombinantly expressed double-acetylated NPM1 was performed by incubating the lysate with 50 μL of Pierce^TM^ Anti-*c*-Myc Magnetic Beads (Thermo Scientific) for 1 h at room temperature. Beads were pre-washed with 1X TBS-T buffer (25 mM Tris, 150 mM NaCl, 0.05% Tween-20) prior to incubation. After incubation, the beads were collected with a magnetic stand and washed twice with 5X TBS-T buffer (125 mM Tris, 750 mM NaCl, 0.25% Tween-20), followed by one was with ultrapure water. Protein elution was achieved by adding 100 µL of 100 mM glycine, pH 2.0 and incubating in agitation for 10 min at room temperature. The supernatant was collected and neutralized with 15 μL 1M Tris-HCl pH8.

The sample fractions were resolved in 10% SDS-PAGE gels and then transferred onto Amersham^TM^ Hybond^TM^ PVDF membranes (Cytiva). Membranes were blocked with 5% BSA in TBS-T for 30 min and immunoblotted overnight at 4 °C with mouse anti-*c*-myc tag antibody (1:1000) (05-724, Sigma Aldrich) or mouse anti-acetyl-lysine antibody (1:500) (sc-32268, Santa Cruz Biotechnology). After washing, membranes were incubated for 1 h at room temperature with HRP-conjugated anti-mouse IgG secondary antibody (A9044, Sigma Aldrich). The immunoreactive bands were detected using Immobilon Western Chemiluminescent HRP Substrate (WBKLS0500, Millipore).

## Molecular Dynamics simulations

Molecular dynamics (MD) simulations were carried out using AMBER 14SB force field (Maier *et al*., 2015) in explicit solvent using OPC water models under periodic boundary conditions using an orthorhombic cell geometry, with a minimal distance between protein and cell walls of 10 Å. The net charge of the system was neutralized including chlorine and sodium atoms. The crystallographic structure of the pentameric array of the N-terminal domain of NPM1 (PDB accession number: 4N8M; Mitrea *et al*., 2014) was used as the starting structure of the simulations, and as a template to generate the modified acetylated and acetyl-mimetic variants. Charges and force-field parameters for the acetyl-lysine residue (Papamokos *et al*., 2012) were used to generate coordinates and topology files with AMBER’s LeaP module. All NPM1 variants were capped with an N-methylamide (NHE) group to neutralize the non-native charge of the free C-terminus.

Equilibrium MD simulations were performed using OpenMM (Eastman *et al*., 2017) v7.4.2. Each system was subjected to 1 µs MD simulations at 298 K at a constant pressure of 1 atm after 2,500 steps of energy minimization, using Particle mesh Ewald (PME) electrostatics with an Ewald summation cut-off of 9 Å to account for Coulomb and van der Waals interactions. Replicates (n=3) were run using varying initial coordinates and topology parameters. The simulation’s trajectory statistics were analyzed using the CPPTRAJ (Roe *et al*., 2013) module of AMBER. Mdtraj (McGibbon *et al*., 2015) Python module was used to compute the most frequent residue pair contacts (0.35 nm cutoff) involving positions 27 or 54 in each system. For each residue pair, the mean contact frequency between monomers was considered.

## Protein expression and purification

For protein expression of WT and Lys-to-Gln/Arg variants of NPM1, *E. coli* BL21(DE3) (70235, Novagen) cells were transformed with the respective pET28a(+) plasmid and grown at 37 °C overnight on liquid lysogeny broth (LB) medium supplemented with 50 µg mL^-1^ kanamycin, that was then used to inoculate 1 L cultures in 2 L flasks (1:10 dilution). Protein expression was induced at an optical density (OD600) of 0.6-0.8 by adding 1 mM isopropyl-β-D-thiogalactoside (IPTG) (R3679, iNtRON Biotechnology). Cultures were grown for 20 h at 30 °C and 180 rpm. For the expression of C*c*, *E. coli* BL21 (DE3) cells were grown in LB medium supplemented with 100 µg mL^-1^ ampicillin for 20 h at 30 °C and 180 rpm.

For C*c* protein expression, *E. coli* BL21 (DE3) cells transformed with the pBTR1 plasmid were cultured in LB supplemented with 100 μg mL^-1^ ampicillin for 20 h at 30 °C and 180 rpm. The p19ARF_(1-37)_ synthetic peptide MGRRFLVTVRIQRAGRPLQERVFLVKFVRSRRPRTAS was purchased from GeneCust (France). The peptide was weighted and reconstituted in the appropriate buffer without any observed aggregation.

After NPM1 protein expression, cells were collected at 4,500 *g* for 10 min and resuspended in a lysis buffer composed of 20 mM Tris-HCl (pH 8.0), 800 mM NaCl, 10 mM Imidazole, 1 mM phenylmethylsulfonyl fluoride (PMSF), cOmplete^TM^ EDTA-free Protease Inhibitor Cocktail (Roche), 0.2 mg mL^-1^ lysozyme and 0.02 mg mL^-1^ DNaseI (Roche). The cell extract was sonicated for 5 min and centrifuged at 27000 *g* for 1 h to remove cell debris. Protein purification of NPM1 was carried out by affinity chromatography in a column with Ni-NTA resin (HisPur^TM^ Ni-NTA Resin, Thermo Scientific^TM^) previously equilibrated with lysis buffer. Protein was eluted using a 10-300 mM imidazole step gradient. Protein-containing elution fractions were dialyzed in 10 mM sodium phosphate buffer (pH 7.4) and concentrated in Amicon® Ultra-15 Centrifugal Filter Units (10 kDa cutoff) until reaching the required concentration. Sample purity and protein integrity was assessed by SDS-PAGE and protein concentration was quantified through Bradford protein assay. For C*c* purification, cells were resuspended in 10 mM tricine buffer pH 8.5 supplemented with cOmplete^TM^ EDTA-free Protease Inhibitor Cocktail, 0.2 mg mL^-1^ lysozyme and 0.02 mg mL^-1^ DNaseI. After sonication and centrifugation, the cell lysate was subjected to cation exchange chromatography as previously reported (Gomila *et al*., 2022). Fractions containing pure C*c* were dialyzed against 10 mM sodium phosphate, pH 7.4, and concentrated in an Amicon® Ultra-15 Centrifugal Filter Unit (3 kDa cutoff). Protein concentration was determined by UV/Vis spectrophotometry, using an extinction coefficient of 28.92 mM^-1^ cm^-1^.

## SDS-PAGE

1 µg of each pure NPM1 protein variant prepared in 10 mM sodium phosphate, 150 mM NaCl, pH 7.4 buffer was mixed with reducing Laemmli SDS sample buffer, heated at the specified temperature for 5 min and separated in 10% SDS-PAGE gels at 200V and RT. Gels were stained with Coomassie Blue.

## Fluorescent Labeling

Cysteine residues in the NPM1 protein variants were labeled using the Oregon Green™ 488 maleimide reagent (Molecular Probes). C*c* and p19ARF_1-37_ were labeled with Alexa Fluor™ 594 NHS Ester (A20004, Invitrogen) by means of their free primary amines under pH 7.4, favoring reaction at the N-terminus. In both cases, a 10-fold molar excess of the dye was mixed with the protein from a concentrated DMSO stock, and the reaction was incubated for 1 h at room temperature in the dark, in a buffer containing 10 mM sodium phosphate and 150 mM NaCl at pH 7.4. The unbound dye was subsequently removed by extensive dialysis against 10 mM sodium phosphate, 150 mM NaCl, and 2 mM DTT, pH 7.4. Protein concentration and labeling efficiency were assessed by UV-visible spectroscopy according to the manufacturer’s guidelines.

## Phase Separation Assays

To perform the phase separation assays, the respective solutions were prepared from a concentrated protein stock containing a mix of unlabeled NPM1 with 10% of fluorescently labeled protein in 10 mM sodium phosphate, 150 mM NaCl, 2 mM DTT, pH 7.4 buffer. For homotypic NPM1 phase separation, the NPM1 protein solution was diluted at the respective concentrations in the same buffer and 15% Ficoll^TM^ PM70 (Cytiva) was added. To perform heterotypic phase separation of NPM1-ARF or NPM1-C*c*, the NPM1 protein solution was mixed with the p19ARF_(1-37)_ peptide or C*c* at the indicated concentrations, also in 10 mM sodium phosphate, 150 mM NaCl, 2 mM DTT, pH 7.4 buffer.

For all the preparations, 2 μL of the mixtures were placed on sealed chambers composed by a glass slide, a layer of pierced 100-μm thick double-sided 3M^TM^ tape and a coverslip, as previously described (Casado-Combreras *et al*., 2025; Giner-Arroyo *et al*. 2026). After a 15 min incubation at RT, phase separation was examined using a Leica DM6000B fluorescence microscope (Leica Microsystems), with a 40× objective and an ORCA-ER camera (Hamamatsu). Differential interference contrast (DIC) and fluorescence filter FITCL5 (excitation band-pass, 480/40; emission band-pass, 527/30) were used to visualize Oregon Green^TM^ 488. 10 microscopy field images from two mounted samples were acquired for each experimental condition. LASX (Leica Microsystems) and FIJI (ImageJ) softwares were employed for image processing (Schneider *et al*., 2012) and FIJI’s trainable Weka segmentation tool was used for image segmentation and area occupancy quantification (Arganda-Carreras *et al*., 2017). Mann-Whitney U test were performed in Origin 2020b (OriginLab Corp.) for the statistical analysis of droplet area occupancy.

## Acknowledgements

This work was supported by the Spanish National Research Agency (AEI), Spanish Ministry of Science, Innovation and Universities (MCIU) and ‘ERDF A way of making Europe’ [MCIU/AEI/10.13039/501100011033, grant numbers PID2021-126663NB-I00, PID2024-157414NB-I00, PRE2019-091404 to R.L.G.A. FPU24/00691 to P.R.G.]; Ramon Areces Foundation [LEUCYTO-FRA2025-MAR] and Instituto de Salud Carlos III (ISCIII) [DTS25/00058]. The authors would like to thank Prof. Eyal Arbely for kindly providing the plasmids for incorporation of acetyl-lysine, as well as the Microscopy, Chromatography, Imaging and Centrifuges Services at the Institute of Plant Biochemistry and Photosynthesis (IBVF, Seville) and the Biomolecular Interaction Platform at cicCartuja (Seville).

## Competing interests

The authors declare no competing interests.

## Notes

### Competing Interest Statement

The authors have declared no competing interest.

