## Supporting Information Appendix for "Acetylation facilitates NPM1 anchoring in the nucleolus by boosting liquid–liquid phase separation"

**This file includes:**

Figures S1-S5
Tables S1-S3

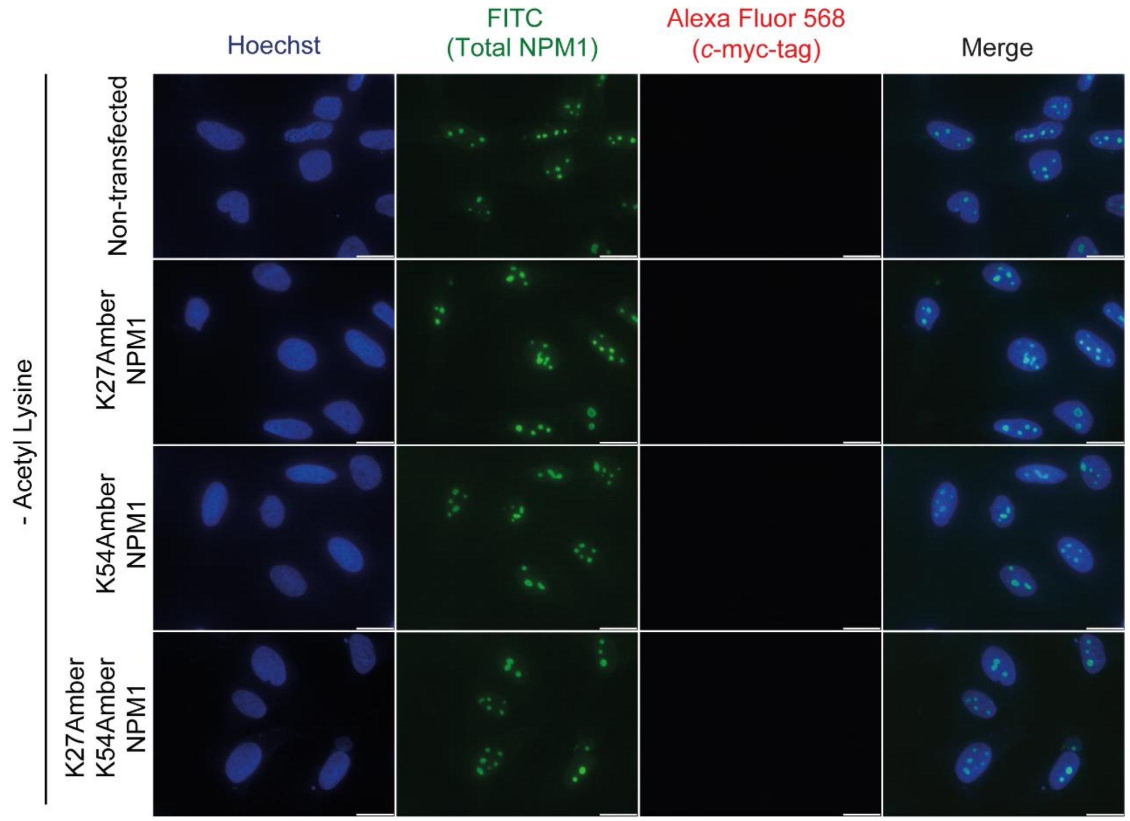


**Figure S1. Fluorescence microscopy images of NPM1 Amber mutants without the addition of AcK.** Immunofluorescence staining of U2OS cells transfected with NPM1 Amber variants without the addition of AcK in the medium. Total and transfected NPM1 were stained using anti-NPM1 and anti-myc tag antibodies, respectively. Scale bars, 20 µm.

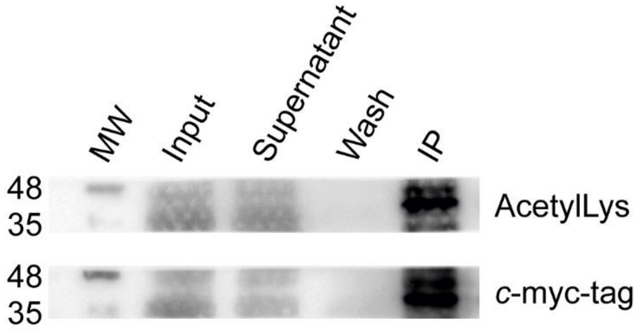


**Figure S2. Immunoprecipitation of *c*-myc-tagged NPM1 K27AcK-K54AcK expressed in U2OS cells.** *c*-myc-tagged NPM1 K27AcK-K54AcK was expressed by genetically encoded AcK incorporation in U2OS cells and immunoprecipitated using anti-*c*-Myc magnetic beads, followed by immunoblotting with anti-acetyl-lysine or anti-*c*-myc-tag antibodies.

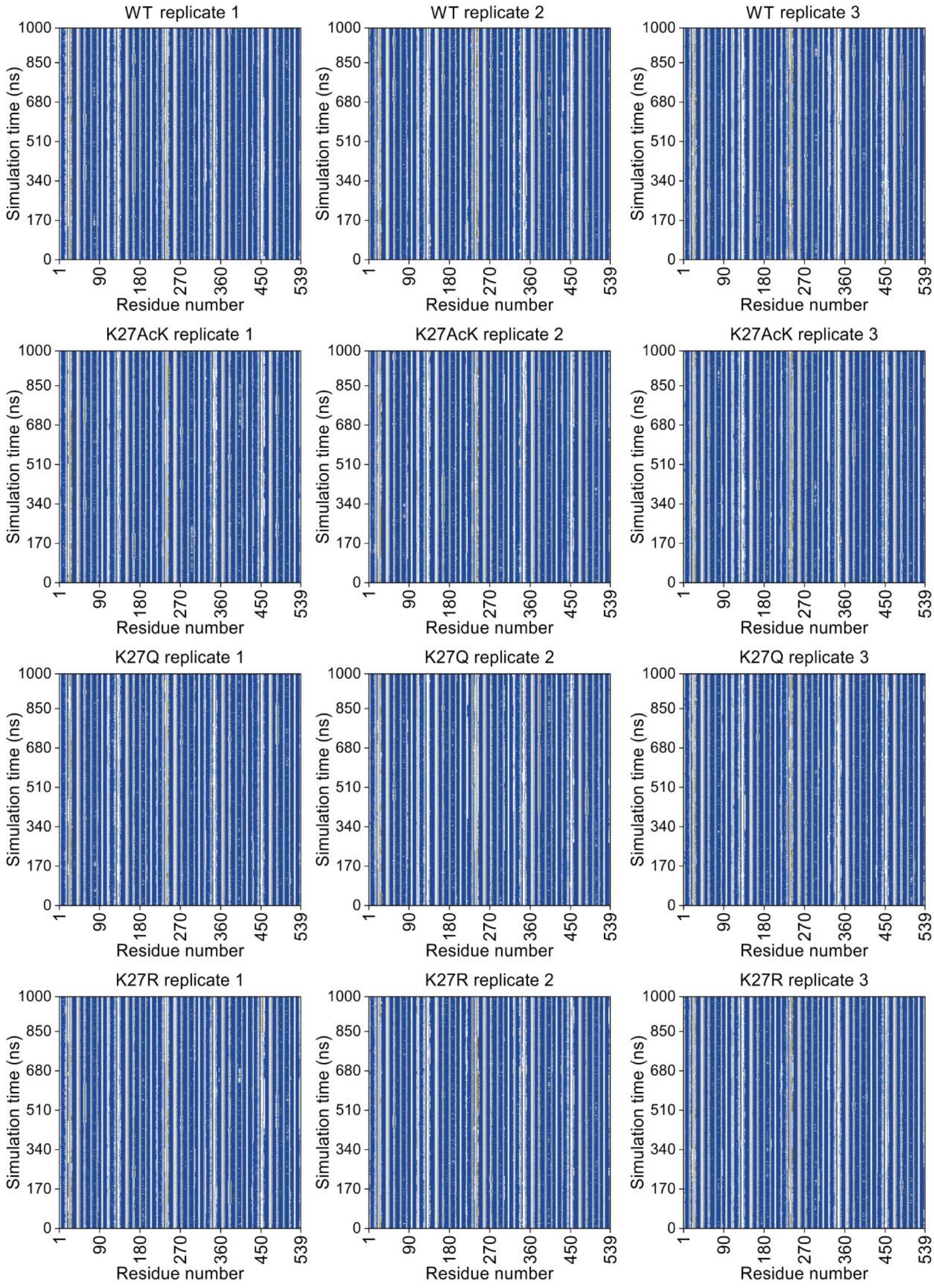


**Figure S3.** Secondary structure Kabsch plots of the three MD simulations replicates of WT and Lys27 mutant NPM1 variants.

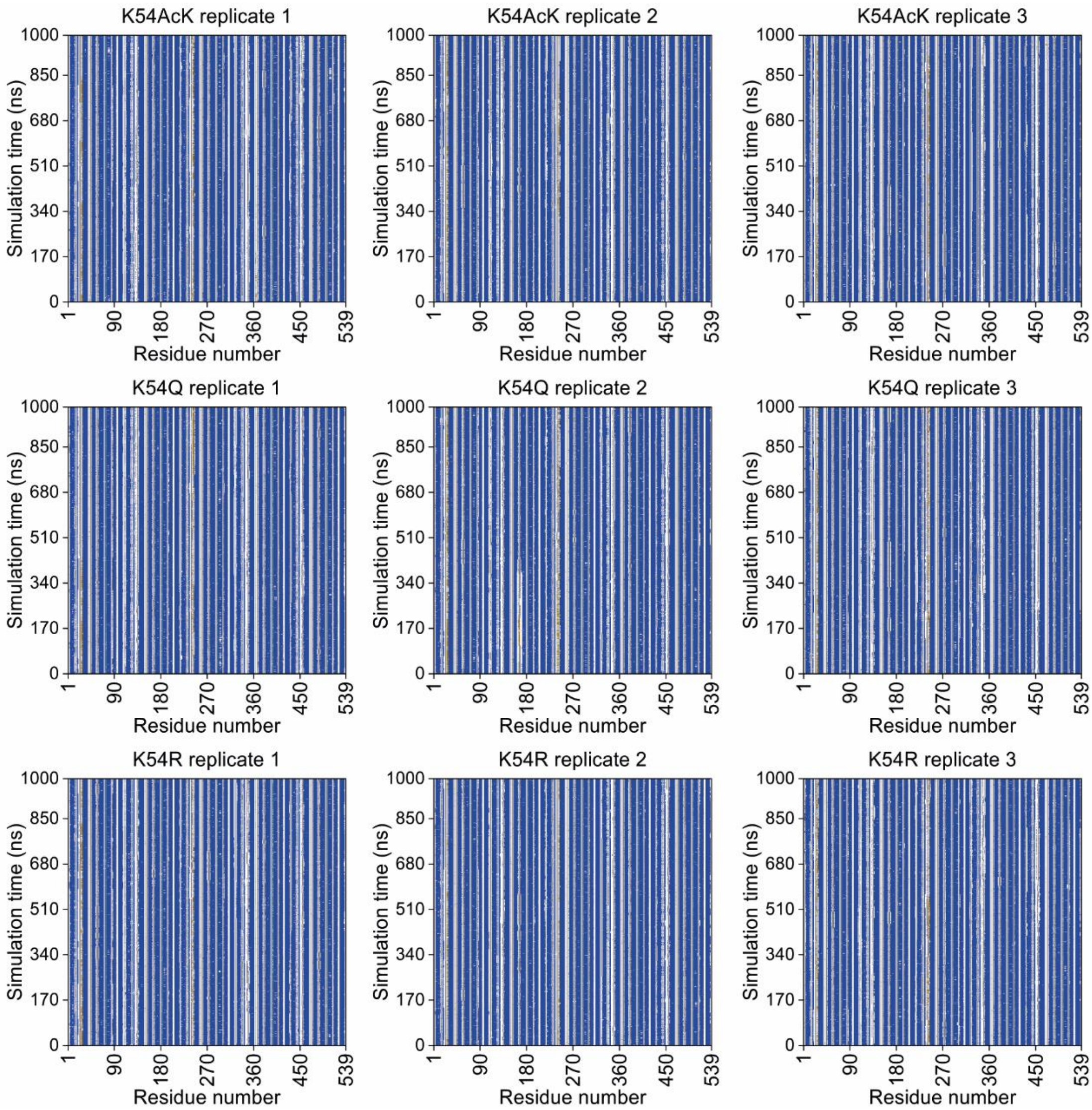


**Figure S4.** Secondary structure Kabsch plots of the three MD simulations replicates of Lys54 mutant NPM1 variants.

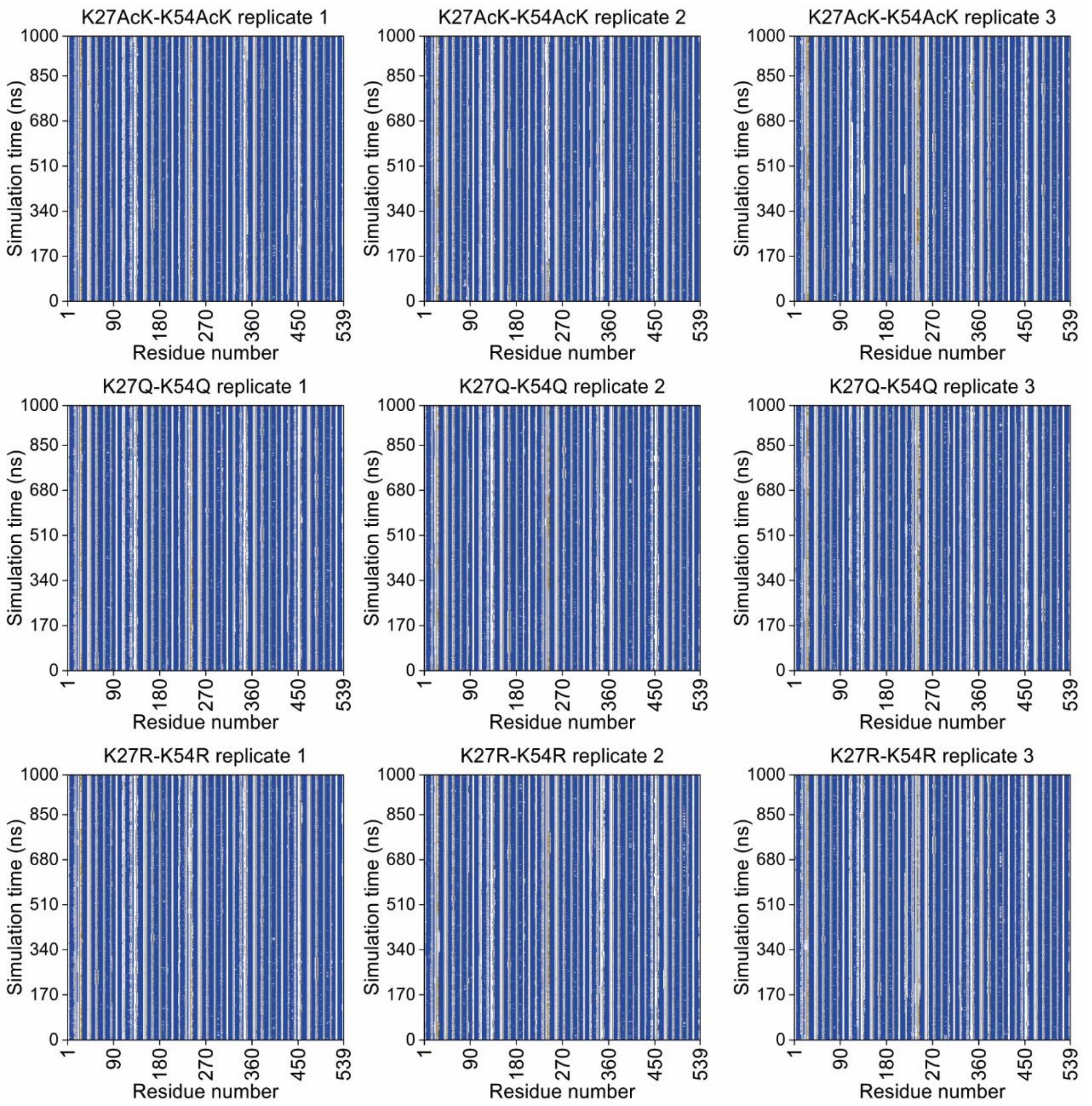


**Figure S5.** Secondary structure Kabsch plots of the three MD simulations replicates of Lys27-Lys54 mutant NPM1 variants.

**Table S1. Oligonucleotides used in this work.**

| Oligonucleotide | Sequence (5'-3') |
|---|---|
| NPM1-K27Amber-Fw | ACTAAAGGCCGACTAGGATTATCACTTT |
| NPM1-K27Amber-Rv | AAAGTGATAATCCTAGTCGGCCTTTAGT |
| NPM1-K54Amber-Fw | GCATAGGATGAGTTGCACATTGTTGAAGCA |
| NPM1-K54Amber-Rv | CATCCTATGCACCAGCCCCTAAACTGAC |
| NPM1-K27Q-Fw | ACTAAAGGCCGACCAAGATTATCACTTT |
| NPM1-K27Q-Rv | AAAGTGATAATCTTGGTCGGCCTTTAGT |
| NPM1-K54Q-Fw | GCACAGGATGAGTTGCACATTGTTGAAGCA |
| NPM1-K54Q-Rv | CATCCTGTGCACCAGCCCCTAAACTGAC |
| NPM1-K27R-Fw | ACTAAAGGCCGACCGAGATTATCACTTT |
| NPM1-K27R-Rv | AAAGTGATAATCTCGGTCGGCCTTTAGT |
| NPM1-K54R-Fw | GCACGTGATGAGTTGCACATTGTTGAAGCA |
| NPM1-K54R-Rv | CATCACGTGCACCAGCCCCTAAACTGAC |
| pBud-NPM1-cloning-Fw | GAAACTCATCTCTGAAGAGGATCTGTAACTCGAGAGATCTGGCCG |
| pBud-NPM1-cloning-Rv | TCCATGTCCATCGAATCTTCCATGGTACCTTCGAAGCGGCC |

**Table S2. Plasmids used in this work.**

| Plasmid | Description | Source |
|---|---|---|
| pBud-AcKRS1-eGFP150Amber-4×tRNA | pBudCE4.1-based plasmid with AcKRS1 tRNA synthetase under CMV promoter, four copies of the pyrolysyl-tRNA synthetase gene, and eGFP150Amber under the EF1a promoter. | (Cohen and Arbely, 2016) |
| pBud-AcKRS1-NPM1-4×tRNA | Vector derived from pBud-AcKRS1-eGFP150TAG-4×tRNA for the mammalian expression of cmyc-tagged NPM1 WT | This study |
| pBud-AcKRS1-NPM1-K27Amber-4×tRNA | Vector derived from pBud-AcKRS1-NPM1-4×tRNA for the mammalian expression of cmyc-tagged NPM1 with genetically encoded acetyl-lysine in position 27 | This study |
| pBud-AcKRS1-NPM1-K54Amber-4×tRNA | Vector derived from pBud-AcKRS1-NPM1-4×tRNA for the mammalian expression of cmyc-tagged NPM1 with genetically encoded acetyl-lysine in position 54 | This study |
| pBud-AcKRS1-NPM1-K27AmberK54Amber-4×tRNA | Vector derived from pBud-AcKRS1-NPM1-4×tRNA for the mammalian expression of cmyc-tagged NPM1 with genetically encoded acetyl-lysine in positions 27 and 54 | This study |
| pcDNA3.1 NPM1-cmyc WT | Mammalian expression vector for cmyc-tagged NPM1 | (Giner-Arroyo *et al.*, 2026) |
| pcDNA3.1 NPM1-cmyc K27Q-K54Q | Mammalian expression vector for cmyc-tagged NPM1 K27Q-K54Q | This study |
| pcDNA3.1 NPM1-cmyc K27R-K54R | Mammalian expression vector for cmyc-tagged NPM1 K27R-K54R | This study |
| pEt28a(+) NPM1 | Bacterial expression vector for NPM1 | (González-Arzola *et al.*, 2022) |
| pEt28a(+) NPM1 K27Q | Bacterial expression vector for NPM1 K27Q | This study |
| pEt28a(+) NPM1 K54Q | Bacterial expression vector for NPM1 K54Q | This study |
| pEt28a(+) NPM1 K27Q-K54Q | Bacterial expression vector for NPM1 K27Q-K54Q | This study |

**Table S3. Solvent exposure of residues in NPM1 oligomerization domain.** The solvent accessibility of the residues in the NPM1 oligomerization domain structure (PDB ID: 4N8M) was calculated using PDBePISA (Krissinel and Henrick, 2007). Inaccessible residues (dark blue), solvent-accessible residues (light blue) and interfacing residues (yellow) are indicated. Residues making hydrogen/disulphide bond, salt bridge or covalent link (HSDC), accessible surface area (ASA, Å²), buried surface area (BSA, Å²), buried area percentage (||||, one bar per 10%) and solvation energy effect (ΔiG, kcal/mol) are reported.

<table>
<tr><th>Residue</th><th>HSDC</th><th>ASA</th><th>BSA</th><th>ΔiG</th></tr>
<tr><td>PRO 14</td><td></td><td>143.11</td><td>0.00</td><td>0.00</td></tr>
<tr><td>GLN 15</td><td></td><td>84.96</td><td>3.88 |</td><td>-0.07</td></tr>
<tr><td>ASN 16</td><td>H</td><td>94.25</td><td>72.06 ||||||||</td><td>-0.55</td></tr>
<tr><td>TYR 17</td><td></td><td>110.01</td><td>59.70 ||||||</td><td>0.88</td></tr>
<tr><td>LEU 18</td><td>H</td><td>164.58</td><td>159.43 ||||||||||</td><td>1.78</td></tr>
<tr><td>PHE 19</td><td></td><td>16.57</td><td>5.84 ||||</td><td>0.05</td></tr>
<tr><td>GLY 20</td><td></td><td>46.87</td><td>25.27 ||||||</td><td>0.18</td></tr>
<tr><td>CYS 21</td><td></td><td>14.45</td><td>0.00</td><td>0.00</td></tr>
<tr><td>GLU 22</td><td></td><td>53.18</td><td>0.00</td><td>0.00</td></tr>
<tr><td>LEU 23</td><td></td><td>0.00</td><td>0.00</td><td>0.00</td></tr>
<tr><td>LYS 24</td><td></td><td>77.73</td><td>0.00</td><td>0.00</td></tr>
<tr><td>ALA 25</td><td></td><td>59.78</td><td>0.00</td><td>0.00</td></tr>
<tr><td>ASP 26</td><td></td><td>121.32</td><td>0.00</td><td>0.00</td></tr>
<tr><td>LYS 27</td><td></td><td>92.83</td><td>0.00</td><td>0.00</td></tr>
<tr><td>ASP 28</td><td></td><td>56.10</td><td>0.00</td><td>0.00</td></tr>
<tr><td>TYR 29</td><td></td><td>73.38</td><td>0.00</td><td>0.00</td></tr>
<tr><td>HIS 30</td><td></td><td>93.30</td><td>0.00</td><td>0.00</td></tr>
<tr><td>PHE 31</td><td></td><td>15.13</td><td>0.00</td><td>0.00</td></tr>
<tr><td>LYS 32</td><td></td><td>143.95</td><td>0.00</td><td>0.00</td></tr>
<tr><td>VAL 33</td><td></td><td>95.29</td><td>0.00</td><td>0.00</td></tr>
<tr><td>ASP 34</td><td></td><td>53.69</td><td>11.93 |||</td><td>-0.14</td></tr>
<tr><td>ASN 35</td><td></td><td>164.42</td><td>0.00</td><td>0.00</td></tr>
<tr><td>ASP 36</td><td>H</td><td>66.02</td><td>11.80 ||</td><td>-0.17</td></tr>
<tr><td>GLU 37</td><td></td><td>181.59</td><td>0.00</td><td>0.00</td></tr>
<tr><td>ASN 38</td><td>H</td><td>62.30</td><td>9.61 ||</td><td>-0.11</td></tr>
<tr><td>GLU 39</td><td></td><td>126.85</td><td>0.00</td><td>0.00</td></tr>
<tr><td>HIS 40</td><td></td><td>24.88</td><td>5.00 |||</td><td>0.08</td></tr>
<tr><td>GLN 41</td><td></td><td>49.42</td><td>0.00</td><td>0.00</td></tr>
<tr><td>LEU 42</td><td></td><td>0.00</td><td>0.00</td><td>0.00</td></tr>
<tr><td>SER 43</td><td></td><td>3.75</td><td>0.00</td><td>0.00</td></tr>
<tr><td>LEU 44</td><td></td><td>0.00</td><td>0.00</td><td>0.00</td></tr>
<tr><td>ARG 45</td><td>H</td><td>99.84</td><td>74.50 ||||||||</td><td>-0.22</td></tr>
<tr><td>THR 46</td><td>H</td><td>40.73</td><td>40.33 ||||||||||</td><td>0.39</td></tr>
<tr><td>VAL 47</td><td></td><td>0.00</td><td>0.00</td><td>0.00</td></tr>
<tr><td>SER 48</td><td></td><td>29.07</td><td>28.66 ||||||||||</td><td>0.21</td></tr>
<tr><td>LEU 49</td><td>H</td><td>19.34</td><td>19.34 ||||||||||</td><td>-0.22</td></tr>
<tr><td>GLY 50</td><td></td><td>5.75</td><td>5.25 ||||||||||</td><td>0.08</td></tr>
<tr><td>ALA 51</td><td></td><td>104.15</td><td>51.63 |||||</td><td>0.62</td></tr>
<tr><td>GLY 52</td><td></td><td>55.87</td><td>1.43 |</td><td>-0.02</td></tr>
<tr><td>ALA 53</td><td></td><td>10.18</td><td>0.00</td><td>0.00</td></tr>
<tr><td>LYS 54</td><td></td><td>142.07</td><td>0.00</td><td>0.00</td></tr>
<tr><td>ASP 55</td><td></td><td>67.03</td><td>0.00</td><td>0.00</td></tr>
<tr><td>GLU 56</td><td></td><td>59.79</td><td>0.00</td><td>0.00</td></tr>
<tr><td>LEU 57</td><td></td><td>86.25</td><td>0.00</td><td>0.00</td></tr>
<tr><td>HIS 58</td><td></td><td>0.00</td><td>0.00</td><td>0.00</td></tr>
<tr><td>ILE 59</td><td></td><td>32.15</td><td>0.00</td><td>0.00</td></tr>
<tr><td>VAL 60</td><td></td><td>0.00</td><td>0.00</td><td>0.00</td></tr>
<tr><td>GLU 61</td><td></td><td>14.95</td><td>0.00</td><td>0.00</td></tr>
<tr><td>ALA 62</td><td></td><td>2.75</td><td>0.00</td><td>0.00</td></tr>
<tr><td>GLU 63</td><td></td><td>32.95</td><td>0.00</td><td>0.00</td></tr>
<tr><td>ALA 64</td><td></td><td>16.41</td><td>0.00</td><td>0.00</td></tr>
<tr><td>MET 65</td><td></td><td>112.97</td><td>0.00</td><td>0.00</td></tr>
<tr><td>ASN 66</td><td></td><td>36.78</td><td>0.00</td><td>0.00</td></tr>
</table>

<table>
<tr><th>Residue</th><th>HSDC</th><th>ASA</th><th>BSA</th><th>ΔiG</th></tr>
<tr><td>TYR 67</td><td></td><td>218.57</td><td>0.00</td><td>0.00</td></tr>
<tr><td>GLU 68</td><td></td><td>150.40</td><td>0.00</td><td>0.00</td></tr>
<tr><td>GLY 69</td><td></td><td>51.83</td><td>0.00</td><td>0.00</td></tr>
<tr><td>SER 70</td><td></td><td>49.52</td><td>0.00</td><td>0.00</td></tr>
<tr><td>PRO 71</td><td></td><td>78.14</td><td>0.00</td><td>0.00</td></tr>
<tr><td>ILE 72</td><td></td><td>73.36</td><td>0.00</td><td>0.00</td></tr>
<tr><td>LYS 73</td><td></td><td>130.50</td><td>0.00</td><td>0.00</td></tr>
<tr><td>VAL 74</td><td></td><td>57.37</td><td>0.00</td><td>0.00</td></tr>
<tr><td>THR 75</td><td></td><td>83.91</td><td>0.00</td><td>0.00</td></tr>
<tr><td>LEU 76</td><td></td><td>17.09</td><td>0.00</td><td>0.00</td></tr>
<tr><td>ALA 77</td><td></td><td>7.19</td><td>0.00</td><td>0.00</td></tr>
<tr><td>THR 78</td><td></td><td>63.47</td><td>0.00</td><td>0.00</td></tr>
<tr><td>LEU 79</td><td></td><td>0.97</td><td>0.00</td><td>0.00</td></tr>
<tr><td>LYS 80</td><td></td><td>65.22</td><td>0.00</td><td>0.00</td></tr>
<tr><td>MET 81</td><td></td><td>70.08</td><td>45.17 |||||||</td><td>0.92</td></tr>
<tr><td>SER 82</td><td></td><td>82.28</td><td>20.79 |||</td><td>0.02</td></tr>
<tr><td>VAL 83</td><td></td><td>115.17</td><td>14.88 ||</td><td>-0.08</td></tr>
<tr><td>GLN 84</td><td></td><td>86.15</td><td>7.62 |</td><td>0.08</td></tr>
<tr><td>PRO 85</td><td></td><td>59.53</td><td>59.53 ||||||||||</td><td>0.95</td></tr>
<tr><td>THR 86</td><td>H</td><td>66.69</td><td>65.54 ||||||||||</td><td>0.22</td></tr>
<tr><td>VAL 87</td><td></td><td>25.34</td><td>0.00</td><td>0.00</td></tr>
<tr><td>SER 88</td><td></td><td>85.58</td><td>40.90 |||||</td><td>0.14</td></tr>
<tr><td>LEU 89</td><td></td><td>41.34</td><td>0.00</td><td>0.00</td></tr>
<tr><td>GLY 90</td><td></td><td>52.42</td><td>6.27 ||</td><td>0.10</td></tr>
<tr><td>GLY 91</td><td></td><td>22.89</td><td>0.00</td><td>0.00</td></tr>
<tr><td>PHE 92</td><td></td><td>88.46</td><td>0.00</td><td>0.00</td></tr>
<tr><td>GLU 93</td><td></td><td>96.13</td><td>0.00</td><td>0.00</td></tr>
<tr><td>ILE 94</td><td></td><td>21.41</td><td>0.00</td><td>0.00</td></tr>
<tr><td>THR 95</td><td></td><td>82.53</td><td>0.00</td><td>0.00</td></tr>
<tr><td>PRO 96</td><td></td><td>16.97</td><td>0.00</td><td>0.00</td></tr>
<tr><td>PRO 97</td><td></td><td>59.49</td><td>0.00</td><td>0.00</td></tr>
<tr><td>VAL 98</td><td></td><td>0.64</td><td>0.00</td><td>0.00</td></tr>
<tr><td>VAL 99</td><td></td><td>24.08</td><td>0.00</td><td>0.00</td></tr>
<tr><td>LEU 100</td><td></td><td>0.17</td><td>0.00</td><td>0.00</td></tr>
<tr><td>ARG 101</td><td></td><td>62.89</td><td>0.00</td><td>0.00</td></tr>
<tr><td>LEU 102</td><td></td><td>18.92</td><td>0.00</td><td>0.00</td></tr>
<tr><td>LYS 103</td><td></td><td>137.16</td><td>0.00</td><td>0.00</td></tr>
<tr><td>CYS 104</td><td></td><td>51.17</td><td>0.00</td><td>0.00</td></tr>
<tr><td>GLY 105</td><td></td><td>26.38</td><td>0.00</td><td>0.00</td></tr>
<tr><td>SER 106</td><td></td><td>57.08</td><td>0.00</td><td>0.00</td></tr>
<tr><td>GLY 107</td><td></td><td>5.62</td><td>0.00</td><td>0.00</td></tr>
<tr><td>PRO 108</td><td></td><td>26.45</td><td>0.00</td><td>0.00</td></tr>
<tr><td>VAL 109</td><td></td><td>0.00</td><td>0.00</td><td>0.00</td></tr>
<tr><td>HIS 110</td><td></td><td>76.34</td><td>56.51 ||||||||</td><td>0.27</td></tr>
<tr><td>ILE 111</td><td></td><td>3.69</td><td>2.96 |||||||||</td><td>-0.02</td></tr>
<tr><td>SER 112</td><td></td><td>40.64</td><td>40.50 ||||||||||</td><td>0.47</td></tr>
<tr><td>GLY 113</td><td></td><td>2.81</td><td>2.81 |||||||||||</td><td>0.03</td></tr>
<tr><td>GLN 114</td><td>H</td><td>45.25</td><td>33.85 ||||||||</td><td>-0.09</td></tr>
<tr><td>HIS 115</td><td></td><td>31.82</td><td>19.50 |||||||</td><td>-0.19</td></tr>
<tr><td>LEU 116</td><td></td><td>38.70</td><td>0.00</td><td>0.00</td></tr>
<tr><td>VAL 117</td><td></td><td>10.54</td><td>10.38 ||||||||||</td><td>0.17</td></tr>
<tr><td>ALA 118</td><td></td><td>42.83</td><td>0.00</td><td>0.00</td></tr>
<tr><td>VAL 119</td><td></td><td>103.86</td><td>0.00</td><td>0.00</td></tr>
</table>
